# Low-protein/high-carbohydrate diet induces AMPK-dependent canonical and non-canonical thermogenic response in subcutaneous adipose tissue

**DOI:** 10.1101/2020.02.26.966002

**Authors:** Katia Aquilano, Francesca Sciarretta, Riccardo Turchi, Bo-Han Li, Marco Rosina, Veronica Ceci, Giulio Guidobaldi, Simona Arena, Chiara D’Ambrosio, Matteo Audano, Illari Salvatori, Barbara Colella, Raffaella Faraonio, Concita Panebianco, Valerio Pazienza, Donatella Caruso, Nico Mitro, Sabrina Di Bartolomeo, Andrea Scaloni, Jing-Ya Li, Daniele Lettieri-Barbato

## Abstract

Low-protein/high-carbohydrate (LPHC) diet promotes metabolic health and longevity in adult humans and animal models. However, the complex molecular underpinnings of how LPHC diet leads to metabolic benefits remain elusive. Through a multi-layered approach, here we observed that LPHC diet promotes an energy-dissipating response consisting in the parallel recruitment of canonical and non-canonical (muscular) thermogenic systems in subcutaneous white adipose tissue (sWAT). In particular, we measured Ucp1 induction in association with up-regulation of actomyosin components and several Serca (Serca1, Serca2a, Serca2b) ATPases. In beige adipocytes, we observed that AMPK activation is responsible for transducing the amino acid lowering in an enhanced fat catabolism, which sustains both Ucp1- and Serca-dependent energy dissipation. Limiting AMPK activation counteracts the expression of brown fat and muscular genes, including Ucp1 and Serca, as well as mitochondrial oxidative genes. We observed that mitochondrial reactive oxygen species are the upstream molecules controlling AMPK-mediated metabolic rewiring in amino acid-restricted beige adipocytes. Our findings delineate a novel metabolic phenotype of responses to amino acid shortage, which recapitulates some of the benefits of cool temperature in sWAT. In conclusion, this highlights LPHC diet as a valuable and practicable strategy to prevent metabolic diseases through the enhancement of mitochondrial oxidative metabolism and the recruitment of different energy dissipating routes in beige adipocytes.

## INTRODUCTION

Adipose tissue dynamically readapts its metabolism and phenotype to environmental perturbations. Beige adipocytes, having intermediate features between white and brown adipocytes counterparts, were identified to occur interspersed mainly in subcutaneous white adipose tissue (sWAT) and to acquire brown-like phenotype upon certain environmental stimuli, such as cold exposure or exercise (Aldiss et al., 2018; Petrovic et al., 2010; Wang and Seale, 2016; Wu et al., 2012). Beige/brown adipocytes express the uncoupling protein 1 (Ucp1), which plays a central role in heat production by decoupling transport of protons across the inner mitochondrial membrane from synthesis of ATP. Ucp1-independent sources of heat production have been also identified in beige/brown adipocytes, such as creatine-dependent cycles and a futile cycle of Ca^2+^ shuttling into and out of the endoplasmic reticulum via the SERCA and ryanodine receptors (Lettieri-Barbato, 2019). Recently, the rearrangement of the adipocyte cytoskeleton towards muscle-like signatures has been demonstrated to be fundamental for sustaining non-shivering thermogenesis. In particular, tensional responses generated by actomyosin are critical for the acute induction of oxidative metabolism and uncoupled respiration (Tharp et al., 2018). Moreover, it has been revealed that a set of beige adipocyte precursors may also differentiate in response to thermal stress through a glycolytic myogenic intermediate that is required for cellular homeostasis and survival (Chen et al., 2019).

When activated, thermogenesis promotes metabolic flexibility mainly catabolizing glucose and fats with a beneficial impact on overall body metabolism (Harms and Seale, 2013). Impinging thermogenesis in WAT to overwhelm metabolic diseases is an emerging and promising field of research (Kaisanlahti and Glumoff, 2019). A strategy to activate thermogenesis in WAT could be exercise or cold exposure, but it would seem difficult to apply this condition in daily life in human. For this reason, a number of attempts have been made to find more applicable strategies for treating such metabolic diseases.

Recent reports have suggested that reactive oxygen species (ROS) play a critical role in promoting thermogenesis in white/beige adipocytes (Chouchani et al., 2016; Lettieri Barbato et al., 2015a). Remarkably, thiols redox perturbations induce brown fat-like changes in sWAT and improve metabolic flexibility through the enhancement of glucose and fatty acid oxidation (FAO) (Lettieri Barbato et al., 2015b). The serine/threonine AMPK is a crucial sensor of both energy and redox state (Coughlan et al., 2013; Lettieri Barbato et al., 2014; Zhao et al., 2017) virtually in all cells and tissues. In adipose cells, AMPK promotes fatty acid oxidation (FAO) and Ucp1-dependent and -independent thermogenesis (Ahmadian et al., 2011; Pollard et al., 2019). AMPK limits obese phenotype promoting Serca1-mediated Ca^2+^cycling that is fueled almost entirely by ATP generated from glucose and FAO (Pollard et al., 2019), thus contributing to the improvement of glucose and lipid clearance in the bloodstream. For these reasons, AMPK currently represents an attractive therapeutic target for age-related metabolic disorders (Day et al., 2017).

Nutrients control a dense network of signaling pathways that converge on metabolism by regulating health status (Fontana et al., 2010; Swinburn et al., 2011). Altering dietary macronutrient composition, while keeping the total number of calories constant, is an intriguing strategy that improves body metabolism (Simpson et al., 2017; Solon-Biet et al., 2015b). In particular, it has been demonstrated that mice consuming a low-protein/high-carbohydrate (LPHC) diet lived longer if compared with normal chow diet-fed mice (Solon-Biet et al., 2014); a randomized controlled trial showed that LPHC diet promotes leanness and decreases fasting blood glucose in humans (Fontana et al., 2016). However, the mechanisms leading to amelioration of metabolic health remain poorly understood and the identification of molecular checkpoints mediating these responses may led to identification of druggable targets to improve systemic metabolism.

Herein, we deciphered the molecular and metabolic responses to LPHC diet showing that, similarly to cold exposure, LPHC diet improves oxidative metabolism and induces brown and muscular-like signatures in sWAT. Through multi-layered approaches, we uncovered that beige adipocytes directly sense the amino acid shortage and activate a redox-sensitive and AMPK-mediated signaling that promotes metabolic rewiring and energy dissipation in beige fat cells.

## RESULTS

### Acquirement of muscular-like signatures accompanies sWAT browning upon LPHC diet

Although LPHC diet was identified as effective in ameliorating systemic metabolism and extend lifespan (Laeger et al., 2014; Le Couteur et al., 2016), a deep systematic investigation to decipher whether WAT could be involved has never been carried out. In this study, adult mice fed with a typical western diet (WD, 23P:57C) were changed to LPHC diet (7P:73C) for 2 weeks. To obtain a preliminary perspective of changes induced by this diet, we used a proteomic approach. A pool of sWAT obtained from six/eight mice/group was analyzed. We found that 75 out of the 2383 detected proteins were differentially represented (FC>1.5), among which Ucp1, Cpt1b, Cox7a, Ndufb11, Slc25a20 resulted amongst the ones showing the highest increased changes (**Fig. 1A**). GO terms for biological processes revealed an enrichment of proteins participating in mitochondrial respiratory chain, tricarboxylic acid cycle (TCA cycle), mitochondrial fatty acid β-oxidation (FAO) and response to cold (**Fig. 1B**). Enrichment analysis using Tissue Atlas depicted a distribution of mapped proteins across brown adipose fat, heart and skeletal muscle following LPHC in sWAT (**Fig. 1C**). Interestingly, 77% of the up-regulated proteins pertained to mitochondrial compartment and included matrix as well as inner/outer membrane proteins (**Fig. 1D**), suggesting a higher mitochondrial abundance in sWAT of LPHC than WD group. Consistent with proteomic findings, immunoblotting analyses confirmed increased levels of mitochondrial matrix (Pdhb), inner (Ndufb8) and outer membrane (Tomm20) proteins (**Fig. 1E**). The analysis of Ucp1 protein confirmed its increase both in total homogenates (**Fig. 1F**) and crude mitochondria of sWAT (**Supplemental Fig. 1A**), whereas no significant modulation in Ucp1 protein levels was observed in BAT (**Fig. 1F** and **Supplemental Fig. 1A**). The Ucp1 changes observed in sWAT of LPHC-fed mice were at a level comparable to those observed upon cold exposure (**Supplemental Fig. 1B**), suggesting the occurrence of brown-like features in sWAT upon LPHC diet. In line with these findings, we found reduced sWAT mass likely due to a reduction of adipocyte size and white-to-brown conversion (**Supplemental Fig. 1C**). A higher oxygen consumption rate in sWAT of LPHC than WD group was also observed through a polarographic method (**Supplemental Fig. 1D**). On the contrary, changes neither in mass nor in oxygen consumption were revealed in BAT (**Supplemental Fig. 1C** and **1D**). In line with the metabolic benefits already described for LPHC diet, a higher glucose tolerance at the later phase of oral glucose load was registered in mice fed with LPHC diet with respect to WD group (**Supplemental Fig. 1E**).

**Figure 1.**
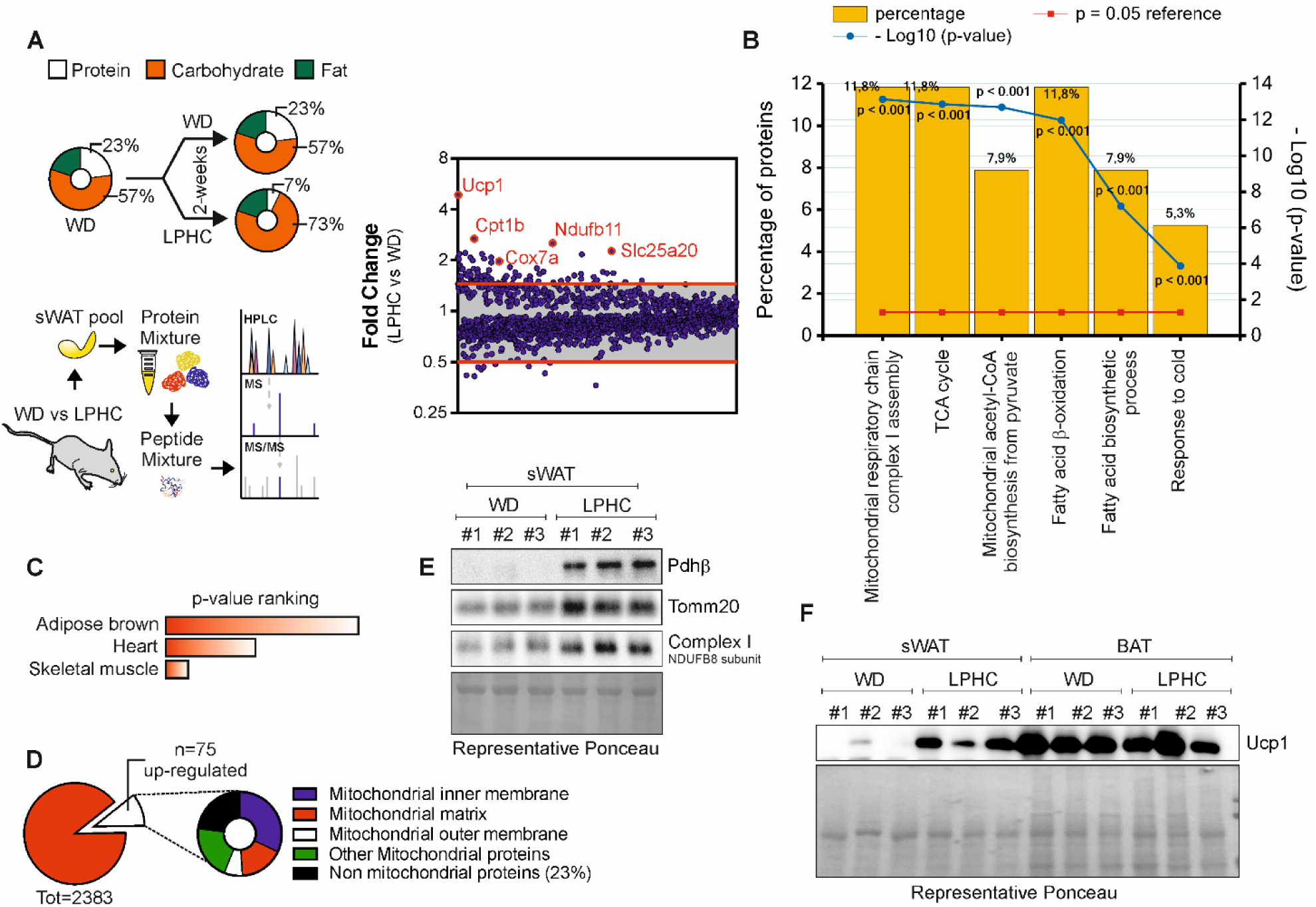
LPHC diet promotes brown fat-like features in sWAT. **A.** Schematic representation of experimental plan and differentially expressed proteins (FC>1.5) from whole proteome profiling in a pool of sWAT isolated from adult male mice fed with WD (n=8 mice) or LPHC (n=6 mice) for 2 weeks. Orange circles were used to mark the most over-represented proteins. **B.** Functional enrichment analysis of over-represented proteins in LPHC fed mice. Gene ontology (GO) terms for biological processes were obtained by FunRich 3.1.3 (http://www.funrich.org). **C.** Over-represented proteins in LPHC fed mice were analysed using Tissue Atlas through the web-based Enrichr bioinformatics tool and sorted by rank based ranking. **D.** Functional enrichment analysis of over-represented proteins in LPHC fed mice. Gene ontology (GO) terms for cellular components were analysed by FunRich 3.1.3 (http://www.funrich.org). **E.** Representative immunoblots of mitochondrial proteins in sWAT of mice fed with WD or LPHC diet for 2 weeks. Ponceau staining was used as loading control. **F.** Representative immunoblots of Ucp1 protein in sWAT and BAT of mice fed with WD or LPHC diet for 2 weeks. Ponceau staining was used as loading control.

We next aimed at better deciphering the responses of sWAT to LPHC diet by performing a deep RNA sequencing; corresponding results were represented by means of a heatmap (**Fig. 2A**). Through pairwise differential gene expression analysis, we found 416 up-regulated (Log_2_FC> 0.58) and 52 down-regulated (Log_2_FC< −0.7) gene transcripts (**Fig. 2B**). The most representative GO terms for biological processes of up-regulated genes (p<0.001) revealed a significant enrichment in the genes of mitochondrial fatty acids catabolism (orange circles) and response to cold (white circle). Notably, an unexpected induction of muscle contraction genes (purple circle) was also found in sWAT of LPHC-fed mice (**Fig. 2C**). Comparative analysis in the abundance of gene transcripts between pWAT and BAT (GSE109829) revealed the overrepresentation of a number of muscular gene transcripts in BAT (**Supplemental Fig. 2A**), arguing that beige adipocytes may also develop non-canonical brown fat genes during thermogenic stimulation. With the aim to identify common putative mechanisms of action between LPHC diet and cold exposure, the up-regulated gene transcripts obtained from sWAT of LPHC-fed mice were integrated with counterparts obtained from sWAT of cold-exposed mice (GSE84860). Notably, the enrichment analysis for biological processes of the 150 overlapping genes pictured a peculiar interaction network among genes involved in cellular respiration, muscle contraction and fatty acid metabolic processes (**Fig. 2D**). A heatmap was depicted, which reports the key common up-regulated genes in sWAT of LPHC fed-mice and cold-exposed mice pertaining to biological processes such as muscle contraction, brown fat, FAO and electron transport chain (**Fig. 2E**). Transcriptomic data were then validated by single gene expression analysis (**Supplemental Fig. 2B**).

**Figure 2.**
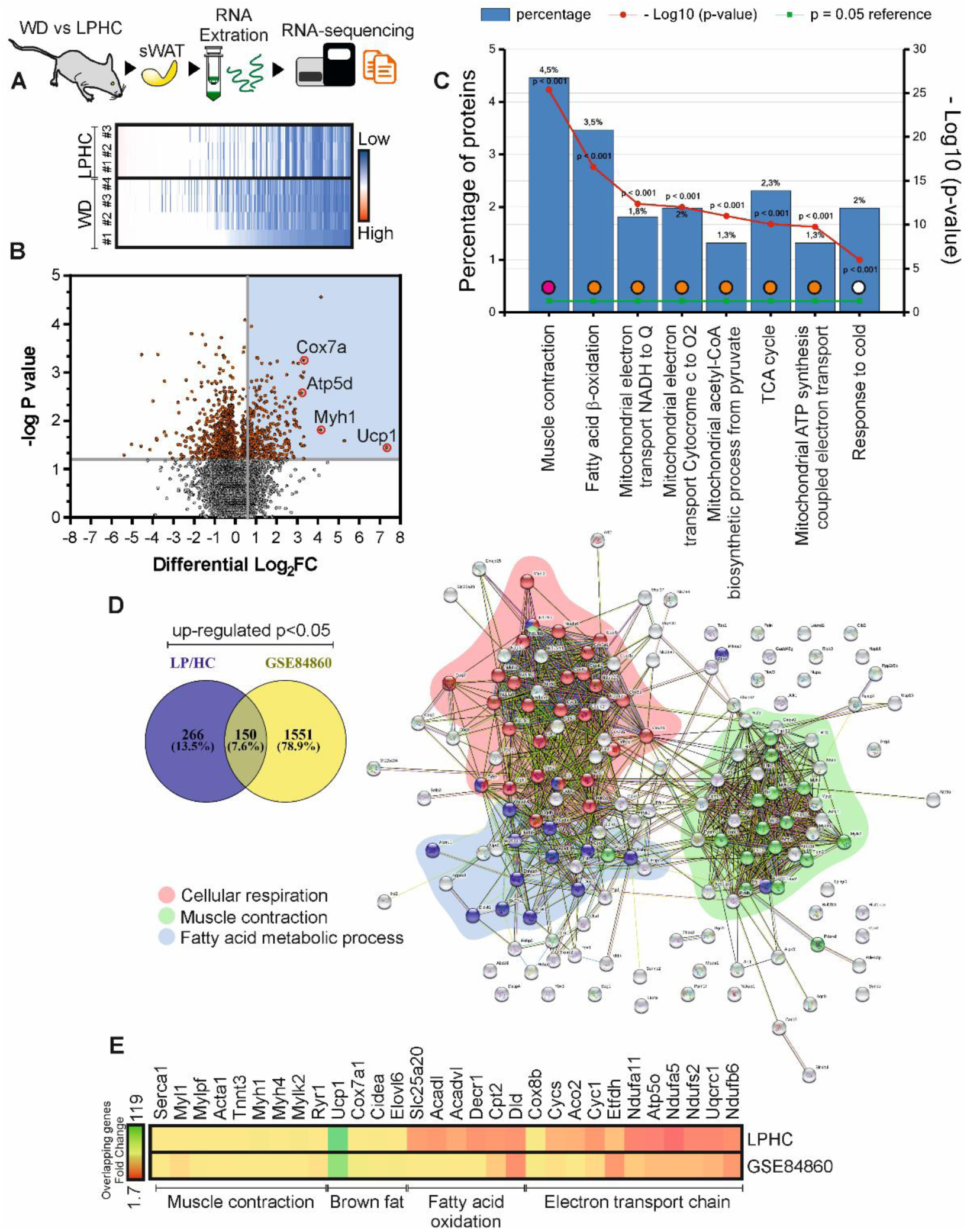
Muscular gene induction accompanies sWAT browning. **A.** Schematic representation of experimental plan and RNA-sequencing of sWAT isolated from adult male mice fed with a WD (n=4 mice) or LPHC diet (n=3 mice) for 2 weeks. Raw RPKM expression values are represented as a heatmap. **B.** Volcano plot representing differentially expressed genes in sWAT isolated from adult male mice fed with LPHC diet (n=3 mice) for 2 weeks versus WD (n=4 mice). Up-regulated genes (Log_2_FC>0.58, p<0.05; n=416, blue area) were reported. **C.** Functional enrichment analysis of up-regulated gene transcripts in LPHC fed mice. Gene ontology (GO) terms for biological processes were analysed by FunRich 3.1.3 (http://www.funrich.org). **D, E**. Venn diagram of up-regulated gene transcripts in sWAT following cold exposure (GSE84860) or LPHC diet. Protein-protein interaction network of overlapped genes (150 genes) was evidenced by STRING (https://string-db.org) setting an interaction score of 0.400. Genes clustered for the main biological processes were reported in the coloured areas (D). Representative common gene transcripts were reported as raw data in a heatmap (E).

### Amino acid lowering is responsible for metabolic rearrangement and sWAT browning

Change in gut microbial composition has been proposed as the mediator of cold-related metabolic benefits (Chevalier et al., 2015). As diet significantly modulates gut microbiota, we performed a metagenomics analysis in order to assess its possible contribution in mediating the effects of LPHC diet on sWAT. As reported in **Supplemental Fig. 3**, a mild microbiota reshaping was induced by LPHC diet; no changes were revealed in the *phyla* mainly affected by cold exposure, such as Verrucomicrobia, Bacteroidetes and Firmicutes (Chevalier et al., 2015), suggesting that the cold-like metabolic benefits of LPHC diet are independent of gut microbiota.

Prior data demonstrated that WAT browning is induced as a response to nutrient shortage (Holmes, 2017). The obtained results opened the question if the reduced amino acid availability was sufficient in triggering the molecular reorganization observed in sWAT following LPHC diet. To explore this issue, we firstly aimed at quantifying the amino acid abundance in sWAT of mice exposed to cool temperature, which is the best-known stimulus promoting white-to-brown conversion of sWAT. PCA highlighted temperature-specific distribution of amino acids (**Fig. 3A**), and the heatmap revealed that exposure to cold significantly decreased the amount of several amino acids in sWAT (**Fig. 3B**). In particular, a diminished concentration of non-polar (glycine, alanine, proline, valine) (**Fig. 3C**), polar uncharged (tyrosine, asparagine, serine, threonine, glutamine) (**Fig. 3D**), positively charged (histidine) (**Fig. 3E**) and negatively charged (aspartate and glutamate) (**Fig. 3F**) amino acids was observed. Reduced amount of citrulline and taurine were also found in sWAT in response to cold (**Fig. 3G**). Such modulation of amino acid abundance was not observed in BAT (**Fig. 3A-G**), which by contrast showed a more proficient reduction of glycolytic and TCA metabolic intermediates than sWAT (**Supplemental Fig. 4A, B**). Actually, glycolysis metabolites remained unchanged in sWAT of cold-exposed mice, while only few derivatives pertaining to the TCA cycle were reduced, i.e. acetyl-CoA, malate and oxaloacetate (**Supplemental Fig. 4A, B**). These responses in association with the significant reduction in the NADPH (**Fig. 3H**), NADH (**Fig. 3I**), ATP and ADP levels (**Fig. 3J**) suggested a strict link between amino acid loss and metabolic rewiring in sWAT.

**Fig. 3.**
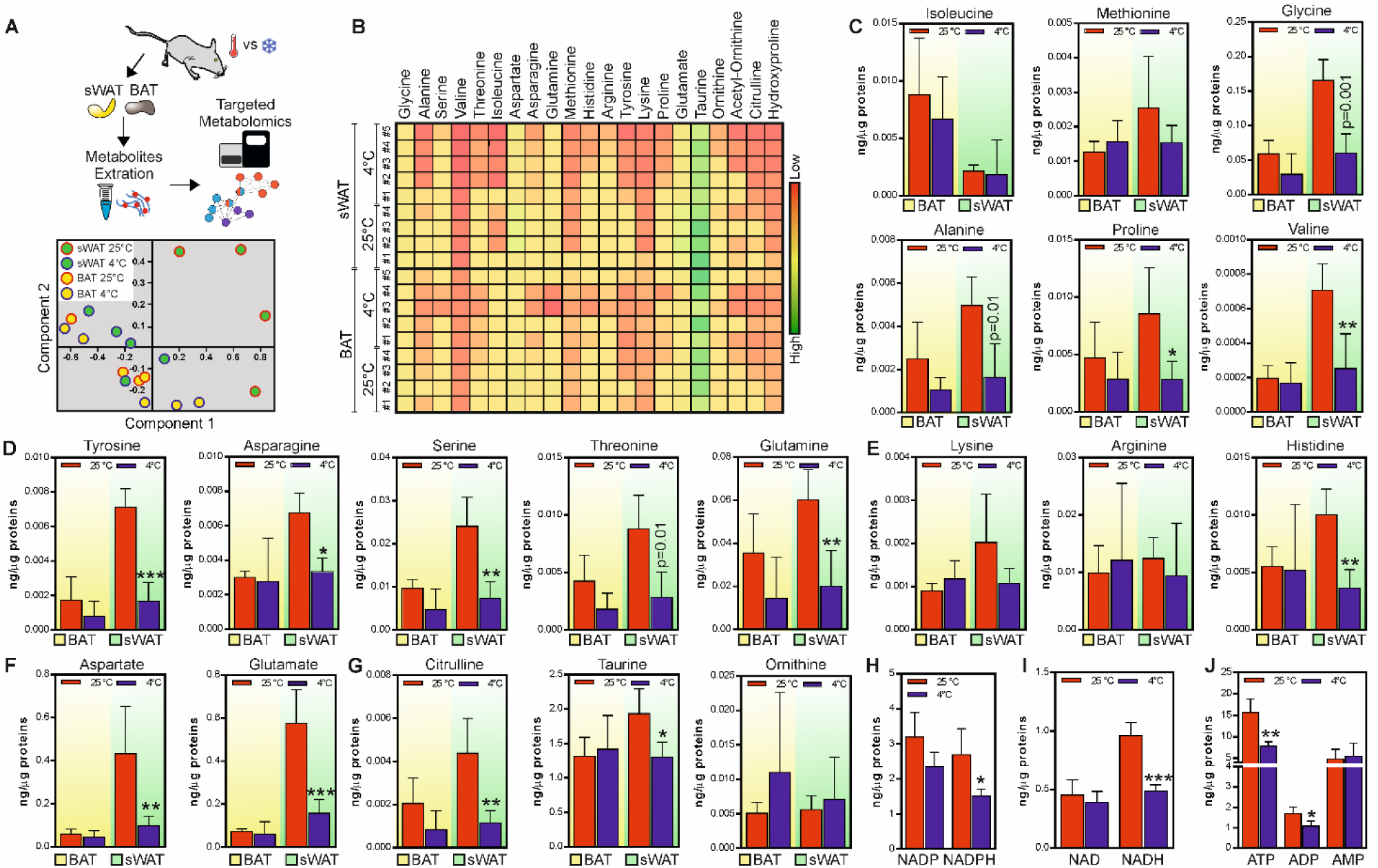
Cool temperature induces amino acid loss in sWAT. **A.** Schematic representation of experimental plan and Principal Component Analysis (PCA) of the metabolites measured in sWAT and BAT of mice exposed to room temperature (25 °C, n=4 mice) or cool temperature (4 °C, n=5 mice) for 24 h. Coloured dots represent individual samples. **B.** Heatmap of biologically major amino acids measured in sWAT and BAT of mice exposed to room temperature (25 °C, n=4 mice) or cool temperature (4 °C, n=5 mice) for 24 h. **C-G.** Single amino acid abundance measured in in sWAT (green area) and BAT (yellow area) of mice exposed to room temperature (25 °C, n=4 mice) or cool temperature (4 °C, n=5 mice) for 24 h. Data are presented as mean ± S.D. *p<0.05; **p<0.01;***p<0.001 25 °C *vs* 4 °C (Student t-test). **H-J.** Metabolites targeting energy metabolism were measured in sWAT (green area) and BAT (yellow area) of mice exposed to room temperature (25 °C, n=4 mice) or cool temperature (4 °C, n=5 mice) for 24 h. Data are presented as mean ± S.D. *p<0.05; **p<0.01;***p<0.001 25 °C *vs* 4 °C (Student t-test).

To investigate whether amino acid lowering was the genuine inducer of sWAT browning upon LPHC diet, we cultured primary beige adipocytes in a medium poorer in amino acids (Amino Acid Restriction, AAR) than control medium, but containing the same concentration of glucose (17.5 mM) and growth factors. In line with data obtained *in vivo* through LPHC diet, AAR induced canonical (Ucp1, Pgc-1α, and Cidea) and non-canonical brown fat markers including actomyosin (Mylpf, Myh3) and Serca (Serca1, Serca2a, Serca2b and Serca3) genes (**Fig. 4A**). Remarkably, AAR also increased the expression of mitochondrial OxPHOS (ATP6, Cox7a, MTCo1) as well as FAO (Slc25a20, Cpt1b) genes; similar results were obtained with the selective β3 receptor agonist CL316,243 (CL) (**Fig. 4A**).

**Fig. 4.**
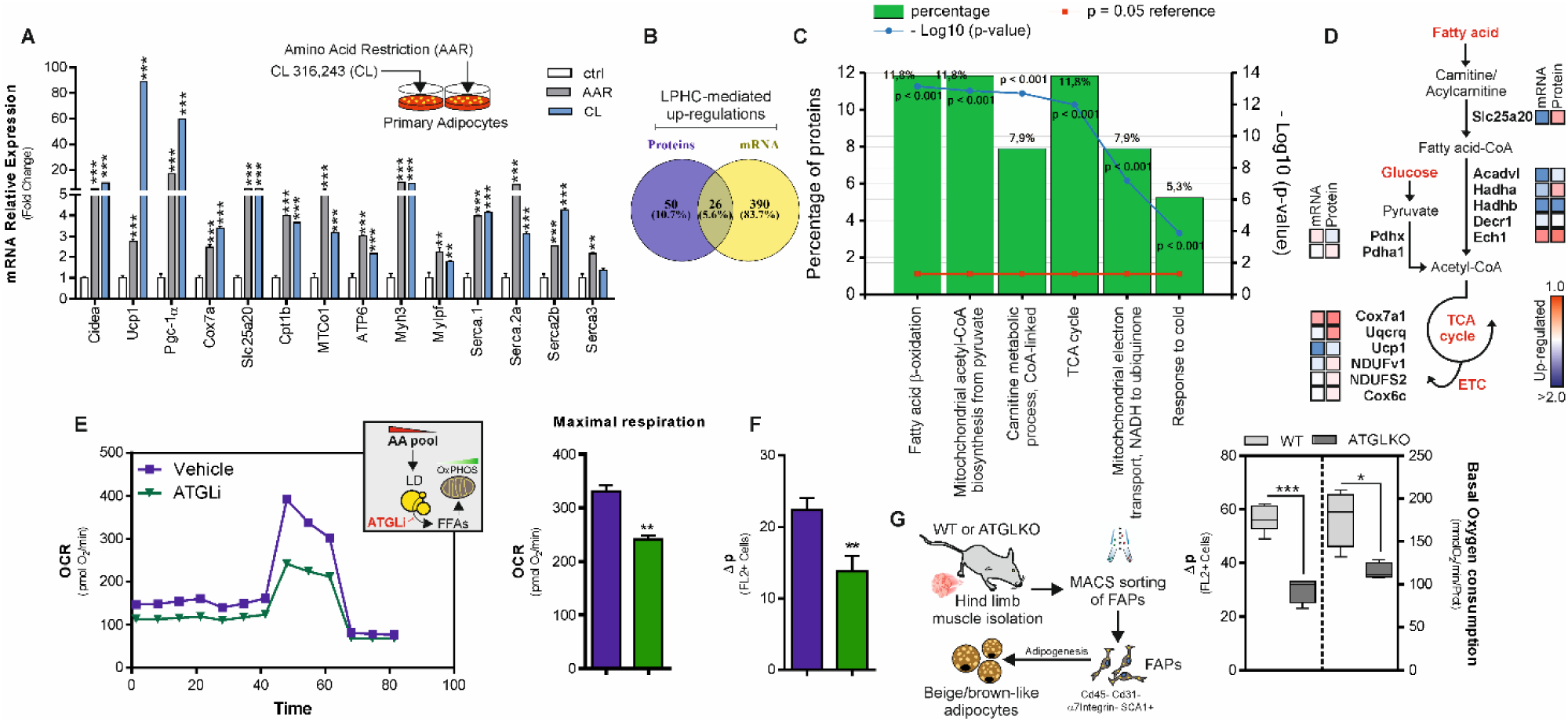
Limiting amino acids enhances Atgl-mediated fatty acid oxidation in beige adipocytes. **A.** Single gene expression analysis in primary beige adipocytes cultured in a medium poor in amino acids (AAR) or adrenergically stimulated by CL316,243 (CL). Data shown are the result of 3 separate experiments. Data are presented as mean ± S.D. ***p<0.001 AAR or CL *vs* Ctr (Student t-test). **B-D.** Venn diagram of up-regulated gene transcripts and proteins in sWAT isolated from adult male mice fed with LPHC diet (n=3 mice) for 2 weeks (B). Functional enrichment analysis of overlapped genes (n=61) was performed by FunRich 3.1.3 (http://www.funrich.org) (C). Schematic representation of key metabolic enzymes over-represented in sWAT (n=61) related to fatty acid oxidation, TCA cycle, ETC and glycolysis pathways. For each enzyme, the corresponding LPHC-induced changes in either mRNA or protein concentration were reported as a heatmap (D). **E.** Seahorse analysis of X9 beige adipocytes cultured in a medium poor in amino acids (AAR) supplemented with Atgl inhibitor Atglistatin (ATGLi) or vehicle. The oxygen consumption rate (OCR) (pmol/min) was monitored for 80 min under basal conditions and upon treatment with the mitochondrial inhibitors oligomycin, FCCP, and rotenone/antimycin. Maximal respiration is reported in the bar graphs. Reported data are the result of 3 separate experiments. Data are presented as mean ± S.E.M. **p<0.01 ATGLi *vs* vehicle (Student t-test). **F.** Mitochondrial proton gradient (Δp) was measured by cytofluorimetry after staining with MitoTracker Red CMXRos in X9 beige adipocytes cultured in a medium poor in amino acids (AAR) supplemented with Atgl inhibitor Atglistatin (ATGLi) or vehicle. Reported data are the result of 3 separate experiments. Data are presented as mean ± S.D. **p<0.01 ATGLi *vs* vehicle (Student t-test). **G.** Schematic representation of FAPs isolated from WT or AtglKO mice. Mitochondrial proton gradient (Δp) and basal oxygen consumption were measured by cytofluorimetry after staining with MitoTracker Red CMXRos and polarography, respectively. Reported data are the result of 3 separate experiments. Data are presented as mean ± S.D. *p<0.05; ***p<0.001 AtglKO *vs* WT (Student t-test).

Overall, these data prompted us to hypothesize that LPHC diet was effective in promoting metabolic rewiring in sWAT. We initially moved at deciphering the metabolic pathway(s) activated by LPHC diet. The up-regulated mRNAs were then integrated with the up-regulated proteins previously detected through transcriptomics and proteomics approaches, respectively (**Fig. 4B**). Among the 61 overlapped genes, we found an enrichment for the biological pathways related to fatty acid catabolism such FAO, TCA cycle and electron transport chain (**Fig. 4C**). The key up-regulated genes pertaining to such biological processes were depicted in a metabolic diagram (**Fig. 4D**).

Our group previously reported that nutrient starved adipocytes catabolize endogenous fatty acids by activating Atgl, the rate-limiting enzyme of lipolysis (Lettieri Barbato et al., 2014). To test whether fat catabolism is enhanced in response to AAR, we inhibited Atgl through the specific inhibitor Atglistatin (ATGLi), and real-time monitoring of cell metabolism was conducted through Seahorse XF analyzer. ATGLi-treated beige adipocytes showed a reduced maximal respiratory capacity (**Fig. 4E**). As expected, ATGLi-treated adipocytes failed to oxidize endogenous fatty acids and became dependent on exogenous fatty acid administration to sustain mitochondrial respiratory capacity (**data not shown**). Consistent with the metabolic measurements, we also found a diminished proton motive force (Δp) in AAR beige adipocytes treated with ATGLi (**Fig. 4F**). The role of Atgl in controlling mitochondrial oxidative metabolism upon AAR was also observed in fibro/adipogenic progenitors (FAPs) isolated from skeletal muscles of WT and ATGL knock-out (ATGL KO) mice. ATGL KO FAPs differentiated in adipocytes showed a lower Δp and basal oxygen consumption than WT FAPs (**Fig. 4G**).

### AMPK coordinates sWAT browning upon amino acid lowering

Activation of AMP-activated kinase (AMPK) increases glucose uptake, FAO and mitochondrial biogenesis in skeletal muscle during contraction (Baldelli et al., 2014; Jager et al., 2007). In white/beige adipocytes, activated AMPK supports thermogenic changes limiting obese phenotype in mice (Gaidhu et al., 2009; Mottillo et al., 2016; Pollard et al., 2019). Of note both muscle contraction and adipose tissue thermogenesis cause energy dissipation involving peculiar proteins such as SERCA and Ucp1 (Bal et al., 2012). According to these findings, we observed a significant phospho-activation of AMPK (Ampk-pT172) in sWAT of LPHC-fed mice, which was comparable to that obtained in cold-exposed mice (**Fig. 5A, B**). Given the recent finding demonstrating that the constitutive AMPK activation in Ucp1-lacking sWAT limits obesity (Ikeda et al., 2017; Pollard et al., 2019), we integrated the first 200 up-regulated genes of sWAT expressing a constitutively-activated AMPK (GSE120429) with the first 200 up-regulated genes of sWAT from LPHC fed mice. Depicting a Venn diagram, we found 48 overlapping genes (**Fig. 5C**) mainly describing skeletal muscle phenotype in sWAT (**Fig. 5D, yellow area**). To confirm the role of AMPK in developing these features, we used adipose tissue-specific AMPKα1/α2 KO mice (referred to as AKO mice), and we found impaired expression of canonical (Pgc-1α, Slc25a20, Cox8b) and non-canonical muscular (Acta1, Myh3, Mylpf, Serca1, Serca2a and Serca2b) brown fat genes in sWAT (**Fig. 5E**). Differently, modulation of muscle-related genes was not found in BAT of cold-exposed AKO mice (**Supplemental Fig. 5A-H**).

**Fig. 5.**
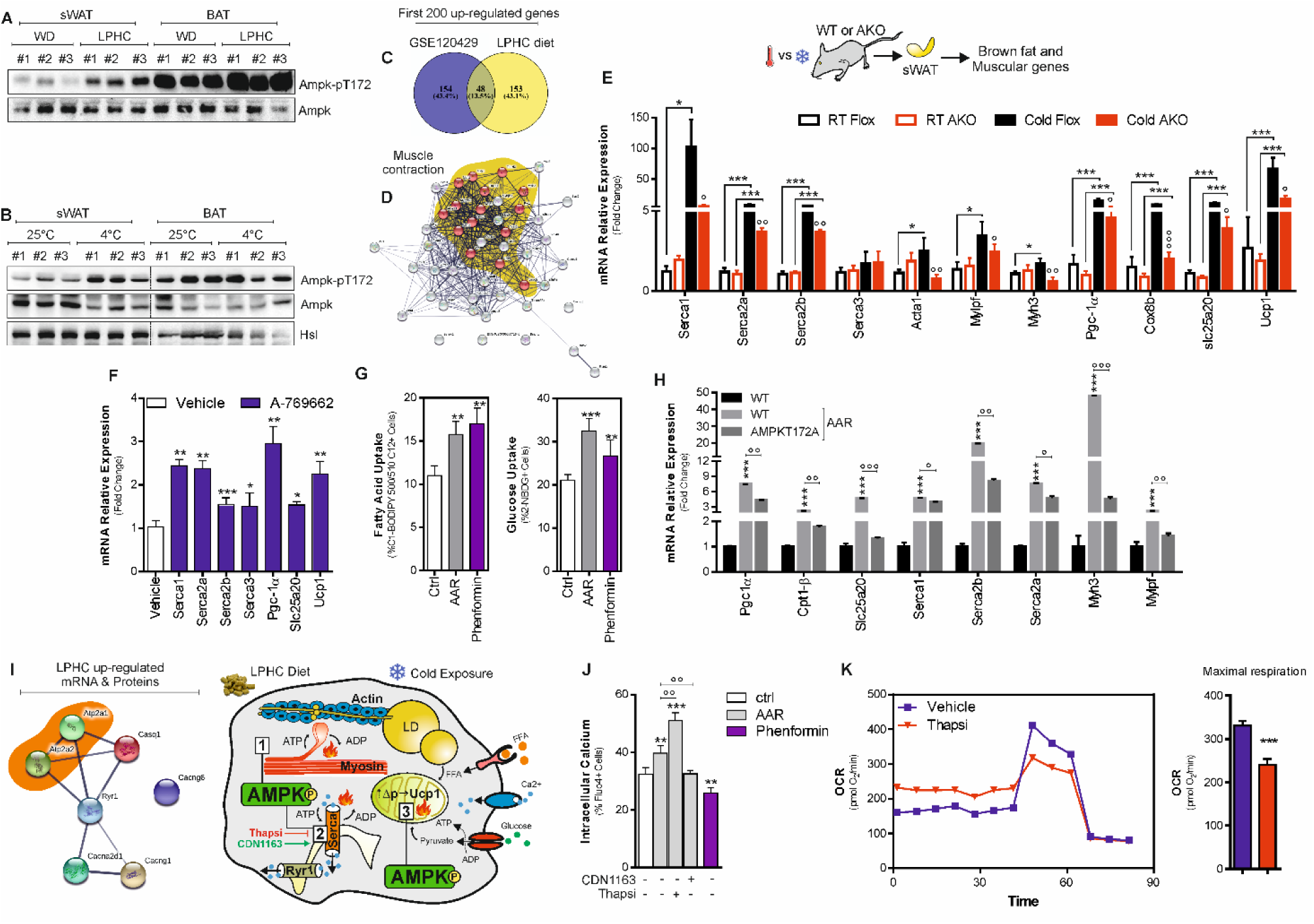
AMPK controls brown fat and muscular genes in sWAT. **A.** Representative immunoblots of AMPK (basal and phosphorylated) in sWAT and BAT of mice fed with WD (n=4 mice) or LPHC diet (n=3 mice) for 2 weeks. **B.** Representative immunoblots of AMPK (basal and phosphorylated) in sWAT and BAT of mice at room temperature (25 °C, n=4 mice) or exposed to cold (4 °C, n=5 mice) for 24 h. Hsl was used as loading control. **C, D**. Venn diagram of the 200 up-regulated gene transcripts obtained from sWAT of mice fed LPHC diet or with constitutively activated AMPK in adipose tissue (GSE120429) (C). Protein-protein interaction network of overlapped genes (n=48) was evidenced by STRING (https://string-db.org) setting an interaction score of 0.400. Yellow area reports the genes clustering muscle contraction Gene Ontology (D). **E.** Single gene expression analysis in sWAT isolated from Flox or AKO mice exposed to room (22 °C) or cool temperature (4 °C) for 48 h. Data are presented as mean ± S.E.M. *p<0.05, ***p<0.001; °p<0.05, °°p<0.01, °°°p<0,001 (one-way ANOVA followed by Dunnett’s multiple comparisons test). **F.** Single gene expression analysis in primary beige adipocytes treated with AMPK agonist (10 µM A-769662). Reported data are the result of 3 separate experiments. Data are presented as mean ± S.E.M. ***p<0.001, **p<0.01, *p<0.05 A-769662 *vs* vehicle (Student t-test). **G.** Glucose and fatty acid uptake were measured in primary beige adipocytes by cytofluorimetry after 2-NBDG or C1-BODIPY 500/510 C12 staining, respectively. Reported data are the result of 3 separate experiments. Data are presented as mean ± S.D. ***p<0.001, **p<0.01 AAR or Phenformin *vs* Ctrl (Student t-test). **H.** Single gene expression analysis in WT and AMPKT172A X9 beige adipocytes cultured in a medium poor in amino acids (AAR). Reported data are the result of 3 separate experiments. Data are presented as mean ± S.D. ***p<0.001 WT AAR *vs* WT; °p<0.05, °°p<0.01, °°°p<0.001 AMPKT172A AAR *vs* WT AAR (one-way ANOVA followed by Dunnett’s multiple comparisons test). **I.** Protein-protein interaction network of up-regulated genes controlling calcium cycle in sWAT following LPHC diet was evidenced by STRING (https://string-db.org) setting an interaction score of 0.400. SERCA proteins were included in the orange area. **J.** Intracellular calcium flux was measured in primary beige adipocytes by cytofluorimetry after staining with Fluo-4, AM probe. Reported data are the result of 3 separate experiments. Data are presented as mean ± S.D. ***p<0.001, **p<0.01 AAR or Phenformin *vs* Ctrl; °° p<0.01 Thapsigargin (Thapsi) or CDN1163 *vs* AAR (one-way ANOVA followed by Dunnett’s multiple comparisons test). **K.** Seahorse analysis of X9 beige adipocytes cultured in a medium poor in amino acids (AAR) treated with the SERCA inhibitor Thapsigargin (Thapsi) or vehicle. The oxygen consumption rate (OCR) (pmol/min) was monitored for 80 min under basal conditions and upon treatment with the mitochondrial inhibitors oligomycin, FCCP, and rotenone/antimycin. Maximal respiration is reported in the bar graphs. Reported data are the result of 3 separate experiments. Data are presented as mean ± S.D. ***p<0.001 Thapsi *vs* vehicle (Student t-test).

Based on these results, we asked whether AMPK activation was directly implicated in the metabolic and molecular rewiring induced by AAR in beige adipocytes. After confirming the occurrence of phospho-activation of AMPK (**Supplemental Fig. 5I**), we analyzed the fatty acid and glucose uptake; and found that it was increased at levels comparable to those observed using the AMPK agonist phenformin (**Fig. 5G**). Next, we limited AMPK activation by transfecting a dominant negative AMPK mutant (AMPKT172A) in beige adipocytes (**Supplemental Fig. 5J**). In AMPKT172A adipocytes, we detected a reduced mRNA expression of Pgc-1α, Cpt1b, Slc25a20, Serca1, Serca2a, Serca2b, Myh3 and Mylpf genes (**Fig. 5H**). To confirm the role of AMPK in promoting the induction of typical brown fat and muscular genes, primary beige cells were treated with the AMPK agonist A769662 and an up-regulation of Serca1, Serca2a, Serca 2b, Pgc-1α, Ucp1 and Slc25a20 genes was observed (**Fig. 5F**).

In brown adipocytes, a reorganization of cytoskeletal proteins occurs upon adrenergic stimulation, which strongly mimics that of cardiomyocytes. Specifically, a PKA-dependent activation of L-type Ca^2+^channels is elicited that facilitates an influx of Ca^2+^, promoting the ATPase activity of the actomyosin complex (Tharp et al., 2018). Interestingly, as reported in the protein interaction network depicted in **Fig. 5I**, besides Serca (Serca1, Serca2a and Serca2b) and actomyosin complex genes, omics data highlighted a significant up-regulation of other genes controlling Ca^2+^cycling (Cacng6, Cacng1, Casq1, Cacna2d1, Ryr1) in sWAT of LPHC fed mice. In accordance with these data, AAR increased intracellular Ca^2+^ levels in beige adipocytes (**Fig. 5J**). Next, we evaluated the direct involvement of SERCAs in the response to AAR. As reported in **Fig. 5J**, SERCA inhibition by thapsigargin (Thapsi) further increased intracellular Ca^2+^ levels, while SERCA activation by CDN1163 reduced intracellular Ca^2+^ levels in AAR-treated adipocytes. Similar results were obtained by using the AMPK agonist phenformin (**Fig. 5J**), highlighting a direct role of AMPK in increasing SERCA-mediated Ca^2+^ cycle in beige adipocytes. As SERCA ATPase, by increasing ADP levels, stimulates mitochondrial respiration in beige adipocytes (Ikeda et al., 2017), we performed Seahorse XF analyses and observed that SERCA inhibition affected maximal mitochondrial respiration (**Fig. 5K**).

### Amino acid lowering triggers the acquirement of the energy-dissipating phenotype in beige adipocytes

Subsequently, we sought to determine what upstream regulators of canonical and non-canonical brown fat genes are induced in AAR-treated adipocytes. Previous results demonstrated that during the thermogenic process, mtROS are produced shifting intracellular redox status of cysteine thiols towards pro-oxidant conditions that are functional in increasing mitochondrial respiration (Chouchani et al., 2016; Lettieri Barbato et al., 2015a; Lettieri Barbato et al., 2015b). In line with this evidence, AAR increases mtROS production in beige adipocytes (**Fig. 6A**). To implicate redox imbalance in the induction of thermogenic response, we carried out supplementation with the antioxidant N-acetyl-L-cysteine (NAC). Expectedly, NAC was able to buffer the AAR-mediated production of mtROS (**Fig. 6A**), and this event was associated with the limited induction of both canonical brown fat (Ucp1, Pgc-1, Cox7a, Slc25a20) and muscular genes (Serca2b, Serca2a, Mylpf, Myh3) (**Fig. 6B**), as well as the down-regulation of Ucp1 protein and mitochondrial complex subunits (**Fig. 6C**). In line with these results, an inhibition of AMPK (**Fig. 6C**) in association with a decrease of fatty acids and glucose uptake, and intracellular Ca^2+^ levels (**Fig. 6D-F**) was observed upon NAC supplementation, thereby highlighting a role of redox imbalance in controlling the responses of beige adipocytes to AAR. Interestingly, AAR promoted an increase of cystine transporter Slc7a11 (**Fig. 6B**), which is implicated in the maintenance of the intracellular thiol pool (Koppula et al., 2018), indicating an attempt of adipocytes to preserve redox homeostasis. Notably, NAC addition inhibited such event (**Fig. 6B**). To confirm the involvement of redox imbalance in the thermogenic commitment in response to AAR we inhibited Slc7a11 by erastin. As reported in **Fig. 6G**, higher levels of Ucp1 protein as well as of actomyosin and SERCA mRNAs were observed in erastin-treated adipocytes compared to controls. Another thiol oxidizing stimulus, i.e. glutathione depletion by its synthesis inhibitor buthionine sulphoximine (BSO), promoted a significant up-regulation of canonical and non-canonical brown fat markers including Ucp1, actomyosin components and SERCA in beige adipocytes as well (**Fig. 6H**).

**Fig. 6.**
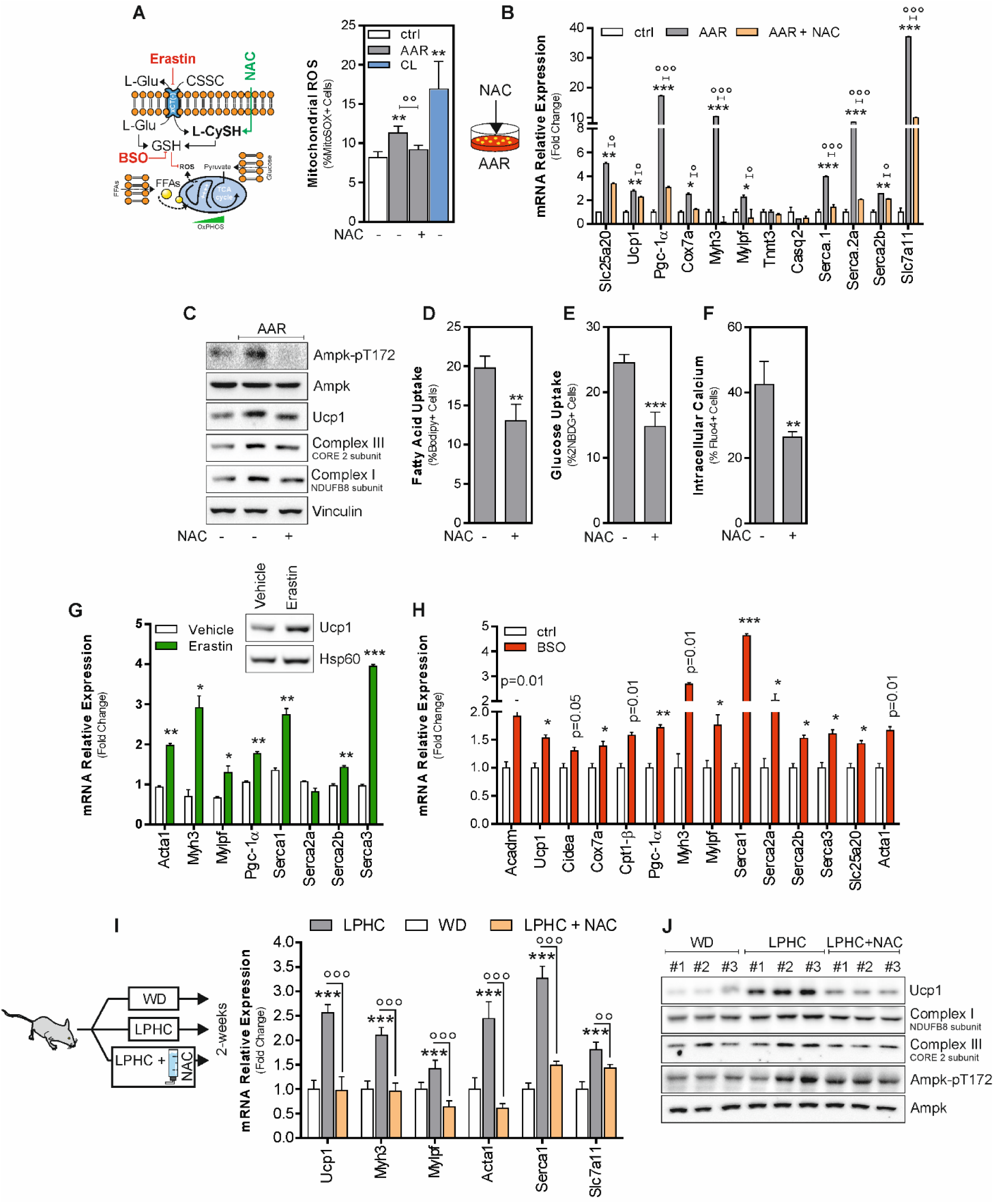
Amino acid restriction perturbs redox homeostasis triggering brown and muscular gene induction in beige adipocytes. **A.** Schematic representation of experimental plan; mitochondrial ROS in X9 beige adipocytes cultured in a medium poor in amino acid (AAR) with NAC or vehicle were measured by cytofluorimetry after MitoSOX staining. Reported data are the result of 3 separate experiments. Data are presented as mean ± S.D. **p<0.01 AAR or CL *vs* control; °° AAR with NAC *vs* AAR (one-way ANOVA followed by Dunnett’s multiple comparisons test). **B.** Single gene expression analysis in X9 beige adipocytes cultured in a medium poor in amino acids (AAR) supplemented with NAC or vehicle. Reported data are the result of 3 separate experiments. Data are presented as mean ± SD. ***p<0.001, **p<0.01 AAR *vs* Ctrl; °p<0.05, °°°p<0.001 AAR with NAC *vs* AAR (one-way ANOVA followed by Dunnett’s multiple comparisons test). **C.** Representative immunoblots of AMPK (basal and phosphorylated), Ucp1, mitochondrial complex III and I subunits in X9 beige adipocytes cultured in a medium poor in amino acids (AAR) supplemented with NAC or vehicle. Vinculin was used as loading control. **D-F**. Fatty acid (D), glucose (E) and calcium (F) uptake in X9 beige adipocytes cultured in a medium poor in amino acids (AAR) supplemented with NAC or vehicle were measured by cytofluorimetry after C1-BODIPY 500/510 C12, 2-NBDG and Fluo-4, AM staining, respectively. Reported data are the result of 3 separate experiments. Data are presented as mean ± S.D. ***p<0.001, **p<0.01 AAR NAC *vs* control (Student t-test). **G, H**. Representative immunoblots of Ucp1 and single gene expression analysis in X9 beige adipocytes treated with Erastin (G) or BSO (H). Hsp60 was used as loading control (G). Reported data are the result of 3 separate experiments. Data are presented as mean ± S.D. ***p<0.001, **p<0.01 *p<0.05 Erastin or BSO *vs* vehicle (Student t-test). **I, J**. Single gene expression analysis (I) and representative immunoblots (J) of AMPK (basal and phosphorylated), Ucp1, mitochondrial complex III and I subunits in sWAT from mice fed WD (n=3), LPHC diet (n=3) or LPHC diet plus NAC. AMPK was used as loading control. Data are presented as mean ± S.D. ***p<0.001, LPHC *vs* WD; °°°p<0.001, °°p<0.01 LPHC plus NAC *vs* LPHC (one-way ANOVA followed by Dunnett’s multiple comparisons test).

Based on these findings, LPHC fed mice were supplemented with NAC by drinking water. Importantly, NAC supplementation also restrained AMPK activation (**Fig. 6J**). As result, diminished levels of Ucp1, subunits of mitochondrial complex I and III, actomyosin components, Serca1 and Slc7a11 were observed in sWAT (**Fig. 6I** and **6J**). Overall above reported findings demonstrate that LPHC diet promotes an acquirement of energy-dissipating signatures in sWAT via an AMPK-dependent redox mechanism.

## DISCUSSION

Mounting evidence suggests that a 1:10 ratio of proteins to carbohydrates protects the body from the damage of aging and extends lifespan. LPHC diet has been demonstrated to reduce age-related neurodegeneration and exert systemic metabolic benefits increasing energy expenditure (Laeger et al., 2014; Solon-Biet et al., 2014; Wahl et al., 2018). WAT takes center place in the control of energy balance and a special emphasis has been given to this tissue as a target of new anti-obesity and anti-diabetic therapeutics (Kaisanlahti and Glumoff, 2019). Although LPHC diet has been proved to ameliorate systemic metabolism in mice and human (Fontana et al., 2016; Levine et al., 2014; Solon-Biet et al., 2015a) and its beneficial effects were at least in part ascribed to the setting of brown-like features in sWAT, the metabolic rearrangement and molecular pathways orchestrating such event are still poorly characterized. Herein, we confirmed the increased glucose tolerance and sWAT browning upon LPHC diet. More importantly, we revealed that LPHC diet promotes the induction of different energy-dissipating routes in sWAT. In the last years, several works gave important proofs supporting that thermogenic adipocytes can use either canonical, *i.e.* Ucp1-mediated thermogenesis, or alternative ways to generate heat, such as creatine cycling and SERCA-mediated Ca^2+^shuttling (Ikeda et al., 2017; Kazak et al., 2015). Ca^2+^ flux from the ER and extracellular medium are actually known responses to adrenergic stimulation in brown adipocytes (Leaver and Pappone, 2002; Lee et al., 1993). Such non-canonical ways contribute to heat production and maintenance of bioenergetics homeostasis by means of exergonic ATP hydrolysis (Ikeda et al., 2017; Pant et al., 2016). Along this line, pharmacological SERCA activation markedly improved glucose tolerance and reduced adipose tissue weight in a murine genetic model of insulin resistance and type 2 diabetes (Kang et al., 2016). To our knowledge, the recruitment of the SERCA in the energy dissipating activity of WAT and BAT has been proposed only as an alternative system to Ucp1 deficiency. Specifically, an evident Serca-dependent Ca^2+^-cycling was identified as a key thermogenic node upon cold exposure in Ucp1-knock out animals (Ikeda et al., 2017; Ukropec et al., 2006). Alternatively, up-regulation of Serca1 and other muscle-like genes was observed in sWAT in a cluster of beige adipocytes that do not express Ucp1 (Pollard et al., 2019).

Our results indicate that, beside the already reported Serca2b system, also other SERCA, namely Serca1 and Serca2a, are up-regulated during sWAT browning; this event occurs in association with Ucp1 up-regulation. Actually, Serca genes were up-regulated concomitant to Ucp1 in AAR-treated adipocytes. The genuine involvement of SERCA-mediated futile Ca^2+^ cycling was demonstrated by the increased intracellular Ca^2+^ levels upon AAR and the ability of pharmacological inhibitor or activator of SERCA to modulate such phenomenon.

More recently, Tharp et al. (Tharp et al., 2018) demonstrated that favoring Ca^2+^-dependent actomyosin mechanics promotes thermogenic capacity of brown adipocytes. Specifically, under this circumstance, the mechanosensitive activation of YAP/TAZ transcription factors is elicited, thereby driving the expression of Ucp1 and other thermogenic genes (Tharp et al., 2018). Notably, we found a significant up-regulation of genes that form the actomyosin complex, arguing that the muscle signatures promoted by LPHC diet and AAR could participate in the thermogenic signaling cascade. It has not to be excluded that being actomyosin head a Ca^2+^-dependent ATP hydrolase, it could be also implicated along with SERCA in the Ucp1-independent energy dissipating activity of sWAT. AMPK is key energy sensor particularly critical in tissues displaying highly changeable energy turnover (Kjobsted et al., 2018). Of note, the expression of skeletal muscle–associated genes was induced in subcutaneous white adipocytes from mice with constitutively induced AMPK (Pollard et al., 2019). Chronic genetic or pharmacological AMPK activation results in protection against diet-induced obesity and induction of energy-dissipating phenotype in white adipose tissue (Pollard et al., 2019; Wu et al., 2018; Zhu et al., 2016). In line with these findings, our experiments carried out in mice with a loss of function of AMPK, specifically in adipose depots (AKO mice), confirmed its role in eliciting the acquirement of canonical and non-canonical(muscular) brown-like signatures in sWAT. Remarkably, amino acid shortage was effective in activating AMPK in white/beige adipocytes, and AMPK mutant adipocytes showed a limited capability to develop muscular and typical brown fat genes. Consistent with these findings, prior reports also showed that AMPK maintains the bioenergetics profile of starved adipocytes by enhancing fatty acid oxidation (Lettieri Barbato et al., 2013). Atgl, which governs the fueling of fatty acids to mitochondrial compartment for degradation (Lettieri Barbato et al., 2014), participates in this signaling process. In line with this assumption, we demonstrated that Atgl inhibition limited respiration activity in AAR-treated adipocytes. Furthermore, fatty acids may also reinforce the Ucp1 uncoupling activity (Fedorenko et al., 2012).

It is worth to notice that mtROS are byproducts of enhanced mitochondrial activity. Several reports demonstrated that mtROS are produced during thermogenesis and are functional in the Ucp1 activation via oxidizing thiols of critical cysteines (Echtay et al., 2002; Lettieri Barbato et al., 2015a; Oelkrug et al., 2014). We found that the development of brown fat-and muscular-like signatures in beige adipocytes upon AAR is associated with mtROS production. Treatment with the antioxidant and thiol reducing agent NAC limited mtROS production, and in turn blunted AMPK activation along with the induction of brown fat and muscular genes in beige adipocytes. Interestingly, in sWAT and beige adipocytes, we found detectable mRNA levels of Slc7a11, a cystine transporter that assures the maintenance of the intracellular thiol redox state. Upon AAR or LPHC diet, Slc7a11 was upregulated, highlighting a cellular attempt to preserve redox homeostasis by maintaining a reduced state. Of notice, oxidation of intracellular thiols is mandatory to sustain thermogenesis, as demonstrated by results showing that GSH decrements is triggered during cold-induced thermogenesis (Lettieri Barbato et al., 2015b), and that chemical inhibition of GSH synthesis by BSO induces sWAT changes that are reminiscent of BAT morphology (Lettieri Barbato et al., 2015b). In line with these findings, through the inhibition of Slc7a11 or GSH synthesis, we were able to recapitulate the responses to LPHC diet. Indeed, through supplementing NAC during LPHC diet, we were able to impede sWAT browning and thermogenesis.

During thermogenesis, adipose depots exploit amino acid, glucose and fatty acid degradation to sustain the uncoupling activity of Ucp1 (Hankir and Klingenspor, 2018; Rose and Richter, 2009; Yoneshiro et al., 2019). Based on our results it can be postulated that changes in the macronutrient ratio, *i.e.* proteins versus carbohydrates or amino acid lowering, enormously increased the energy pressure; this should have committed beige adipose cells to activate both canonical and non-canonical heat producing systems to enhance their energy-dissipating potential. In parallel, amino acid shortage or protein restriction increased metabolic flexibility, as demonstrated by enhanced fatty acid and glucose uptake as well as improved glucose tolerance.

The available data suggest that cold exposure promotes browning program in WAT conferring important systemic metabolic improvements and flexibility in mice and humans. However, such intervention would seem hardly applicable as preventive and therapeutic strategy to increase lifespan and combat age-related metabolic diseases, such as type 2 diabetes. Herein we found that cold exposure causes a significant drop in amino acids amount in sWAT; through reproducing such circumstance by AAR, we were able to induce sWAT browning as well. In conclusion, this study shows that upon LPHC diet, sWAT develops a brown fat and muscular-like phenotype, which presents the recruitment of different canonical and non-canonical energy dissipating routes. The nutrient sensing AMPK governs the induction of Ucp1, SERCA and actomyosin genes via a redox-dependent mechanism. These findings led to suggest that limiting amino acid availability (e.g. via LPHC diet) to beige adipocytes, a metabolic rewiring is induced that enhances metabolic flexibility. Therefore, LPHC diet could represent a more practicable and physiological approach for preventing or correcting the metabolic deregulations associated with age-related diseases.

## METHODS

### Animal husbandry and Treatments

Mouse experimentation was conducted in accordance with accepted standard of humane animal care after the approval by relevant local (The University Animal Welfare Committee – OPBA, Tor Vergata University) and national (Ministry of Health, Legislative Decree No. 26/2014; European Directive 2010/63/UE) committees with authorization n°331/2016-PR and n°324/2018-PR. C57BL/6J adult (3 months-age-old) male mice (purchased from ENVIGO, Italy) were housed one per cage, with a 12-hour light/dark cycle, at 23-25 °C. Mice were weaned with a typical western diet (WD). All animals had free access to water and were randomly assigned to experimental groups. Age and sex-matched mice fed with control (WD) were compared with mice fed with low protein high carbohydrate (LPHC) diet for a total of 3 weeks. Overall analyses were performed after 2 weeks of dietary treatments. Diets were purchased from Research Diets (NJ, USA) and formulated to have the same total energy content (isocaloric) but different ratios in protein to carbohydrate (protein:carbohydrate about 1:10) with fixed fat at 20% of total calories. Each diet was formulated to contain all essential vitamins, minerals, and amino acids for growth in mice. The primary dietary protein component was casein, the main carbohydrate component was starch, and the main fat component was soy oil. NAC was solubilized in drinking water at final concentration of 2 g/L and this solution was refreshed every 48 h. Adult (3 months old) male mice acclimated to room temperature group (25 °C) were compared with cold-exposed counterparts (4 °C), for 24 h. At the end of treatments, blood and adipose tissues were harvested 6 h after fasting to limit confounding factors. Animals were sacrificed in a randomized order to minimize experimental bias, and the adipose tissues were weighed and flash-frozen in liquid nitrogen or fixed for analyses. For LPHC and WD group, each analysis was performed by a pool of n=2 mice per sample (total sample size for WD group n=8 mice; LPHC group n=6 mice).

### Adipose tissue-specific AMPK α1/ α2 double-KO mice

Related animal experiments were approved by the Animal Care and Use Committee of the Shanghai Institute of Materia Medica, where the experiments were conducted. All animals were housed in a temperature-controlled room (22 ± 2 °C), with a light/dark cycle of 12 h. Adipose tissue-specific AMPKα1/α2 double-KO mice (referred to as AKO mice) were generated as previously described (Wu et al., 2018). Male AKO mice and age-matched AMPKα1/α2-floxed littermates were randomly divided into four groups. Cold exposure experiments were performed at 8 weeks of age. AKO or floxed mice were single-caged and exposed to room temperature (22 °C) or cool temperature (4 °C), for 48 h. Food and water were available *ad libitum*. At the end of the study, tissues were dissected, weighed, immediately frozen in liquid nitrogen and stored at −80 °C.

### Oral glucose tolerance test (OGTT)

Oral glucose tolerance test (OGTT) was performed at the end of dietary treatments by oral gavage (2 g of dextrose/kg body mass) in overnight-fasted mice. Blood samples were collected from the tail vein at 0, 15, 30, 60 and 120 min after glucose loading, and the blood glucose level was immediately measured by a commercially available glucometer (Bayer).

### Histochemical analysis

Mouse tissues were fixed in 10% neutral-buffered formalin and embedded in paraffin. Sections (5-μm thick) were stained with hematoxylin and eosin (H&E) according to standard protocols. Microscopy analysis was performed by using a Leica DM6 B microscope at the indicated magnification, and images were captured by a sCMOS camera under the same parameter setting.

### Crude mitochondrial fractions and Oxygen consumption

Crude mitochondria were isolated from sWAT and BAT as previously described(Lettieri Barbato et al., 2015b), and lysed in RIPA buffer (50 mM Tris-HCl, pH 8.0, 150 mM NaCl, 12 mM deoxycholic acid, 0.5% Nonidet P-40, and protease and phosphatase inhibitors) for protein determinations. Oxygen consumption was determined in whole tissues by using the Oxygraph Plus oxygen electrode system (Hansatech Instruments Ltd., Norfolk, UK). In particular, the adipose tissue was maintained in culture medium without FBS and the real-time oxygen consumption rate was normalized for tissue weight.

### RNA-sequencing

Total adipose RNA extraction was isolated with TRIzol Reagent (Invitrogen) and purified using the RNeasy mini kit protocol (Qiagen), according to the manufacturer’s instructions. The collected samples were subject RNA-sequencing using an IlluminaNextSeq500 (Illumina); the indexed libraries were prepared with a TruSeq StrandedmRNA (Illumina) Library Prep kit according to the manufacturer’s instructions. The quality of the single-end reads was evaluated with FastQC v.0.11.5. All resulting files were filtered to remove low quality reads and adapters with Trimmomatic v.0.36 (Bolger et al., 2014). The resulting reads were mapped to the *Mus musculus* genome (GRCm38) with HISAT2 v.2.1.0 (Kim et al., 2015) using default parameters, while Stringtie v1.3.4d (Pertea et al., 2015) was applied to the BAM files obtained with HISAT2 to generate expression estimates and to quantify the transcript abundance as transcripts per kilobase per million of mapped reads (TPM). The count matrices generated by Stringtie were imported in R where differential expression analysis was performed using the Deseq2 package (Love et al., 2014) to compare the two different conditions. The functional annotation was performed through the AnnotationDbi R library. Differential expressed genes were selected with threshold of Log_2_FC>0.58 (*p*<0.05).

### Targeted Metabolomics

For metabolomic analyses, sWAT and BAT were homogenized in 250 µL of methanol/acetonitrile 1:1 (v/v) with D-Glucose-13C6 1 ng/mL (internal standard, Sigma Aldrich, 389374) and centrifuged at 4°C. Supernatant was passed through cellulose filter and saved for subsequent analysis. Amino acid (aa) quantification was performed through previous derivatization. Samples were incubated with PITC (Phenyl isothiocyanate) solution for 20 min at 25°C, dried and resuspended in 5 mM ammonium acetatein MeOH/H2O 1:1 (v/v). Metabolomic data were obtained on an API-4000 triple quadrupole mass spectrometer (AB SCIEX) coupled with a HPLC system (Agilent) and CTC PAL HTS autosampler (PAL System). The identity of all metabolites was confirmed using pure standards. Quantification of different metabolites was performed with a liquid chromatography/tandem mass spectrometry (LC-MS/MS) method using a C18 column (Biocrates) for amino acids and cyano-phase LUNA column (50 mm x 4.6 mm, 5mm; Phenomenex) for the other metabolites. Amino acids were analyzed through a 10 min run in positive while other metabolites were run in negative ion mode in a 5 min run. The mobile phases for positive ion mode analysis (aa) were phase A: 0.2% formic acid in water and phase B: 0.2% formic acid in acetonitrile. The gradient was T0 100% A, T5.5 min 5% A, T7 min 100% A with a flow rate of 500µL/min. The mobile phase for negative ion mode analysis (all other metabolites) was phase A: 5 mM ammonium acetate pH 7.00 in MeOH. The gradient was100% A for all the analysis with a flow rate of 500µL/min. MultiQuant software (version 3.0.2) was used for data analysis and peak review of chromatograms. Quantitative evaluation of all metabolites was performed based on calibration curves with pure standards, then data were normalized on micrograms of proteins.

### 16S rDNA qPCR and Amplicon sequencing

Fecal pellets were immediately frozen in liquid nitrogen after collection and stored at −80 °C. Fecal nucleic acid was extracted from fecal pellets using E.Z.N.A. stool DNA kit (OMEGA, Bio-tek). Bacterial 16S rRNA gene was amplified from total DNA following the Illumina 16S Metagenomic Sequencing Library Preparation instructions. The V3–V4 hypervariable region amplicon was obtained by PCR with universal primers with Illumina adapters (underlined): forward primer: 5′<u>TCGTCGGCAGCGTCAGATGTGTATAAGAGACAG</u>CCTACGGGNGGCWG CAG; reverse primer: 5’<u>GTCTCGTGGGCTCGGAGATGTGTATAAGAGACAG</u>GACTACHVG GGTATCTAATCC selected from Klindworth et al. (Klindworth et al., 2013). Following a first purification step, the V3-V4 amplicon was subjected to a second PCR in order to barcode the libraries using the Illumina dual-index system and then purified again. Next, libraries were diluted to 4nM and pooled before carrying out a paired-end sequencing (2 x 300 cycles) on an Illumina MiSeq device (Illumina Inc., San Diego, CA, USA) according to the manufacturer’s specifications. 16S Metagenomics GAIA 2.0 software (http://www.metagenomics.cloud, Sequentia Biotech 2017; Benchmark of Gaia 2.0 using published datasets available online at: http://gaia.sequentiabiotech.com/benchmark) was used to analyze the Sequence data generated as FASTQ files after performing the quality control of the reads/pairs (i.e. trimming, clipping and adapter removal steps) through FastQC and BBDuk. The reads/pairs are mapped with BWA-MEM against the databases (based on NCBI). The average number of reads per sample was 220,507.1 (SD +/- 104,675.1).

### Cells and Treatments

X9 cells were purchased from ATCC and grown in DMEM/F12 containing 2.5 mM L-glutamine, 15 mM HEPES, 0.5 mM sodium pyruvate, 1200 mg/L sodium bicarbonate, 15%, FBS and L-Alanyl-L-Glutamine final concentration of 2.36 mM. Confluent cells were differentiated to beige adipocytes by DMEM/F-12 containing 5 μM dexamethasone, 0.5 μg/ml insulin, 0.5 mM isobutylmethylxanthine (IBMX), 0.5 μM rosiglitazone, 1 nM Triiodothyronine (T3) and 10% FBS. Three days after induction, cells were maintained in media containing insulin, T3, and 10% FBS until day 8.

For primary beige adipocytes, the stromal-vascular fraction were prepared and differentiated for 8 days, as previously described with some modifications (Lettieri Barbato et al., 2015b). Briefly, inguinal adipose tissue from 6- to 7-week old male C57BL/6J mice was minced on ice and digested with 10 mg/ml collagenase D (Roche) and 2.4 mg/ml dispase II (Roche) in PBS supplemented with 1% bovine serum albumin, for 45 min, at 37 °C. Treatment was quenched by addition of complete medium and products were filtered through a 100 μm strainer (BD Biosciences). The cell suspensions were centrifuged, suspended and filtered through a 40 μm strainer (BD Biosciences), and then further centrifuged and suspended before plating onto 10 cm dishes. SVF cells were cultured in DMEM/F12 supplemented with 10% FBS and 1% penicillin/streptomycin (Invitrogen). Adipocyte differentiation was carried out in growth medium supplemented with 850 nM insulin, 0.5 mM IBMX, 1 μM dexamethasone, 125 nM indomethacin, 1 nM T3 and rosiglitazone 1 μM, for 48 h, and then in growth medium supplemented with 850 nM insulin and 1 μM T3, for additional 6 days.

Fibro/Adipogenic progenitors (FAPs) were isolated from hind limbs of WT and ATGL KO mice and the heterogeneous cell suspension was purified using MACS microbeads technology as previously described (Reggio et al., 2020). For magnetic beads separation, the microbead-conjugated antibodies against CD45 (Cat. No. 130-052-301; Miltenyi Biotech), CD31 (Cat. No. 130-097-418; Miltenyi Biotech) and α7-integrin (Cat. No. 130-104-261; Miltenyi Biotech) were used. FAPs were selected as CD45− CD31− α7-integrin− SCA1+ cells. For adipogenic induction, DMEM supplemented with 20% FBS, 0.5 mM IBMX, 0.4 mM dexamethasone, 1 μg/ml insulin and 1μM rosiglitazone was used. After 48 h, the culture medium was replaced with the maintenance medium consisting of DMEM supplemented with 20% FBS, 1 μg/ml insulin and 1 μM rosiglitazone.

Differentiated adipocytes (day 8) were transfected with a pcDNA3 empty vector or with a pcDNA3 vector containing the Myc-tagged coding sequence for the α2 subunit of AMPK carrying the T→A substitution at residue 172 (AMPKT172A). Amino acid restriction (AAR) was performed with Hank’s Balanced Salt Solution (HBSS) containing the equimolar concentration of glucose to control medium, with 10% FBS and 1% penicillin/streptomycin (Invitrogen), for 4 h. Adrenoceptor stimulation was performed with CL-316,243 (CL) at a final concentrations of 10 μM, for 6 h. N-acetyl cysteine (NAC) and thapsigargin were used at final working concentrations of 5 mM and 2 μM, respectively. Phenformin and CDN1163 were used at final concentrations of 500 μM and 20 μM, respectively, for 6 h. A-769662 was used at final concentrations of 10 μM, for 6 h. Erastin was used in CL-treated adipocytes at final concentration of 10 μM, for 6 h. Buthionine sulfoximine (BSO) was used in serum-free media medium at a final concentration of 1 mM, for 16 h.

### Proteomics analysis

For quantitative proteomic analysis, protein concentration in sWAT samples was determined using the Pierce BCA Protein assay kit™ (Thermo Scientific, Rockford, IL, USA), according to manufacturer’s instructions. An aliquot of each protein sample (100 μg) was adjusted to a final volume of 100 μl with 100 mM TEAB. Proteins were reduced with 5 μl of 200 mM tris (2-carboxyethylphosphine), for 60 min, at 55 °C, and then alkylated by adding 5 μl of 375 mM iodoacetamide in the dark, for 30 min, at 25 °C. Alkylated proteins were then precipitated by addition of 6 vol of cold acetone, pelleted by centrifugation at 8000 x g, for 10 min, at 4 °C, and then air dried. Each sample was digested with freshly prepared trypsin (ratio of enzyme to protein 1:50) in 100 mM TEAB, at 37 °C, overnight. Resulting peptides were labelled with the TMT Label Reagent Set (Thermo-Fisher Scientific, USA), at 25 °C, according to manufacturer’s instructions, using the matching: WD-TMT6-126, LPHC-TMT6-127. After 1 h, reaction was quenched by adding 8 μl of 5% w/v hydroxylamine in each tube, and allowing reaction vortexing for 15 min. For a set of comparative experiments, tagged peptide mixtures were mixed in equal molar ratios (1:1) and vacuum-dried under rotation. Then, pooled TMT-labelled peptide mixtures were suspended in 0.1% trifluoroacetic acid, and fractionated by using the Pierce™High pH Reversed-Phase Peptide fractionation kit (Thermo-Fisher Scientific) to remove unbound TMT reagents, according to manufacturer’s instructions. After fractionation, 8 fractions of TMT-labelled peptides were collected, vacuum-dried and finally reconstituted in 0.1% FA.

TMT-labelled peptide fractions were analyzed on a nanoLC-ESI-Q-Orbitrap-MS/MS platform consisting of an UltiMate 3000 HPLC RSLC nano system (Dionex, USA) coupled to a Q-ExactivePlus mass spectrometer through a Nanoflex ion source (Thermo Fisher Scientific) (Visconti et al., 2019). Peptides were loaded on an Acclaim PepMap TM RSLC C18 column (150 mm × 75 μm ID, 2 μm particles, 100 Å pore size) (Thermo-Fisher Scientific), and eluted with a gradient of solvent B (19.92/80/0.08 v/v/v H2O/ACN/FA) in solvent A (99.9/0.1 v/v H2O/FA), at a flow rate of 300 nl/min. The gradient of solvent B started at 5%, increased to 60% over 125 min, raised to 95% over 1 min, remained at 95% for 8 min, and finally returned to 5% in 1 min, with a column equilibrating step of 20 min before the subsequent chromatographic run. The mass spectrometer operated in data-dependent mode, using a full scan (m/z range 375–1500, nominal resolution of 70,000), followed by MS/MS scans of the 10 most abundant ions. MS/MS spectra were acquired in a scan m/z range 110– 2000, using a normalized collision energy of 32%, an automatic gain control target of 100,000, a maximum ion target of 120 ms, and a resolution of 17,500. A dynamic exclusion of 30 s was used. All MS and MS/MS raw data files per sample were merged for protein identification and relative protein quantification into ProteomeDiscoverervs 2.4 software (Thermo Scientific), enabling the database search by Mascot algorithm v. 2.4.2 (Matrix Science, UK) using the following criteria: UniProtKB protein database (Mus musculus, 17030 protein sequences, 10/2019) including the most common protein contaminants; carbamidomethylation of Cys and TMT6plex modification of lysine and peptide N-terminal as fixed modifications; oxidation of Met, deamidation of Asn and Gln, pyroglutamate formation of Gln, phosphorylation of Ser, Thr and Tyr as variable modifications. Peptide mass tolerance was set to ± 10 ppm and fragment mass tolerance to ± 0.02 Da. Proteolytic enzyme and maximum number of missed cleavages were set to trypsin and 2, respectively. Protein candidates assigned on the basis of at least two sequenced peptides and an individual Mascot Score grater or equal to 25 were considered confidently identified. For quantification, ratios of TMT reporter ion intensities in the MS/MS spectra from raw datasets were used to calculate fold changes between samples. Results were filtered to 1% false discovery rate.

### Seahorse analyses

Fully differentiated adipocytes were plated on SeahorseXFe 96 Microplates (Agilent Technologies) at confluence. After 24 h, the cells were incubated in AAR, for 4 h, before assessing the cellular bioenergetics. Cartridges were hydrated with Seahorse XF Calibrant, overnight, and incubated in the absence of CO_2_, at 37°C. Calibrant was changed and refreshed 1 h before the assay. Mitochondrial stress test was performed according to Agilent’s recommendations. Briefly, the cells were washed four times with Seahorse XF Base Medium supplemented with 10 mM glucose, 1 mM sodium pyruvate, and 2 mM glutamine (pH 7.4 ± 0.01). The cells were incubated in the absence of CO_2_ at 37°C, for 30 min. Mitochondrial inhibitors were sequentially injected at the following final concentrations: 1 μM oligomycin, 1.5 μM carbonyl cyanide-4-(trifluoromethoxy)phenylhydrazone (FCCP), and 1 μM/1 μM rotenone/antimycin (Sigma-Aldrich).

### Immunoblotting

Tissues or cells were lysed in RIPA buffer (50 mM Tris-HCl, pH 8.0, 150 mM NaCl, 12 mM 464 deoxycholic acid, 0.5% Nonidet P-40, and protease and phosphatase inhibitors). Ten μg proteins were loaded on SDS-PAGE and subjected to immunoblotting. Nitrocellulose membranes were incubated with anti-Ampk-p172 (#2531, Cell Signalling Technology), anti-Ucp1 (ab23841, abcam), anti-vDAC (sc-32063, Santa Cruz Biotechnology), anti-Hsp60 (sc-13966, Santa Cruz Biotechnology), anti-Ampk (sc-25792, Santa Cruz Biotechnology), anti-Hsl (#4107, Cell Signalling Technology), Pc (sc-271493, Santa Cruz Biotechnology), anti-Tom20 (sc-11415, Santa Cruz Biotechnolog), anti-Complex I ndufb8 (ms-105, MitoSciences), anti-Complex III Core2 subunit (ms-304, MitoSciences), anti-Vinculin (ab-18058, Abcam), and anti-Pdhβ (gtx-119625, GeneTex) primary antibodies, at 1:1000 dilution. Successively, membranes were incubated with the appropriate horseradish peroxidase-conjugated secondary antibodies. Immunoreactive bands were detected with a FluorChem FC3 System (Protein-Simple, San Jose, CA, USA) after incubation of the membranes with ECL Selected Western Blotting Detection Reagent (GE Healthcare, Pittsburgh, PA, USA). Densitometric analyses of the immunoreactive bands were performed with the FluorChem FC3 Analysis Software.

### Flow cytometer analyses

To monitor intracellular calcium flux, mitochondrial ROS, fatty acids and glucose uptake and mitochondrial proton gradient, FLUO-4 AM, MitoSOX red, BODIPY™ 500/510 C1, C12, 2-NBDG, MitoTracker Red CMXRos probes were used according to manufactures protocols. Flow cytometry analyses were performed by FACScalibur (BD, USA) and analyzed by FlowJo software.

### RT-qPCR gene expression analysis

Total adipose RNA was isolated with TRIzol Reagent (Invitrogen) and purified using the RNeasy mini kit protocol (Qiagen), according to the manufacturer’s instructions. RNA (1 μg) was treated with genomic DNAse and retro-transcripted by using PrimeScriptRT Reagent kit (Takara Bio Inc, Japan). qPCR was performed in triplicate on 50 ng of cDNA, by using validated qPCR primers (BLAST), Applied Biosystems™ Power™ SYBR™ Green Master Mix, and the QuantStudio3 Real-Time PCR System (Thermo Fisher, Whaltam, MA, USA) as previously described (Turchi et al., 2020). The primers used for the RT-qPCR reactions are reported in **Supplemental Table 1**. The reaction was carried out according to the manufacturer’s protocol using QuantStudio 3 Real-Time PCR System; data were analyzed according to 2-ΔΔCt method.

### Bioinformatics and Statistical analyses

Functional enrichment analysis including GO and Kyoto Encyclopedia of Genes and Genomes (KEGG) pathway was performed by using FunRich 3.1.3 (http://www.funrich.org), EnrichR (https://amp.pharm.mssm.edu/Enrichr) or David 6.8 (https://david.ncifcrf.gov). STRING (https://string-db.org) was used to visualize and integrate complex networks of proteomics or transcriptomics data.

Statistical parameters, significance, and the number of animals and *in vitro* experiments were reported in the figure legends. Data were analyzed with Excel and GraphPad Prism 7 (La Jolla, CA, USA). The raw data were analyzed using Student t-test or one-way ANOVA followed by Dunnett’s multiple comparisons test. Data are represented as means ± SEM (for n>3 samples) or means ± SD (for n=3 samples), and *p* values of less than 0.05 were considered statistically significant.

## Data availability

All raw data that support the findings of this study are available from the corresponding author upon reasonable request.

## ACKNOWLEDGMENTS

This work was supported by European Foundation for the Study of Diabetes (EFSD/Lilly 2017), and Italian Ministry of Health (GR-2018-12367588) to D.L.B; by University of Rome Tor Vergata (Uncovering Excellence Program) to K.A.; by the National Natural Science Foundation of China (81470166) to J.Y.L.; by MIUR “Progetto Eccellenza” to Dipartimento di Scienze Farmacologiche e Biomolecolari, Università degli Studi di Milano; by Italian Ministry of Health through Ricerca Corrente 2018-2020 program to V.P and by National Research Council (project Nutrizione, Alimentazione ed Invecchiamento attivo NUTRAGE) to A.S.

## AUTHOR CONTRIBUTION

D.L.B. and K.A. conceptualized, designed the study and wrote the manuscript. F.S., R.T., G.G., V.C., B.C. and S.D.B. performed and analysed *in vivo* and *in vitro* experiments. M.R. isolated FAPs from WT and ATGL KO mice. I.S. and M.R. performed and interpreted data of Seahorse assay. B.H.L and J.Y.L conducted and contributed to the related study with AMPK AKO mice. R.F. performed experiments on AMPKT172A adipocytes. M.A. extracted and analysed sample for metabolomics, M.A., N.M., D.C., analysed and interpreted data of metabolomics analyses. V.P. and C.P. performed and analysed microbiota experiments. S.A., C. D. and A.S. performed, analysed and interpreted proteomics data.

## SUPPLEMENTAL MATERIAL

**Supplemental Table 1.**
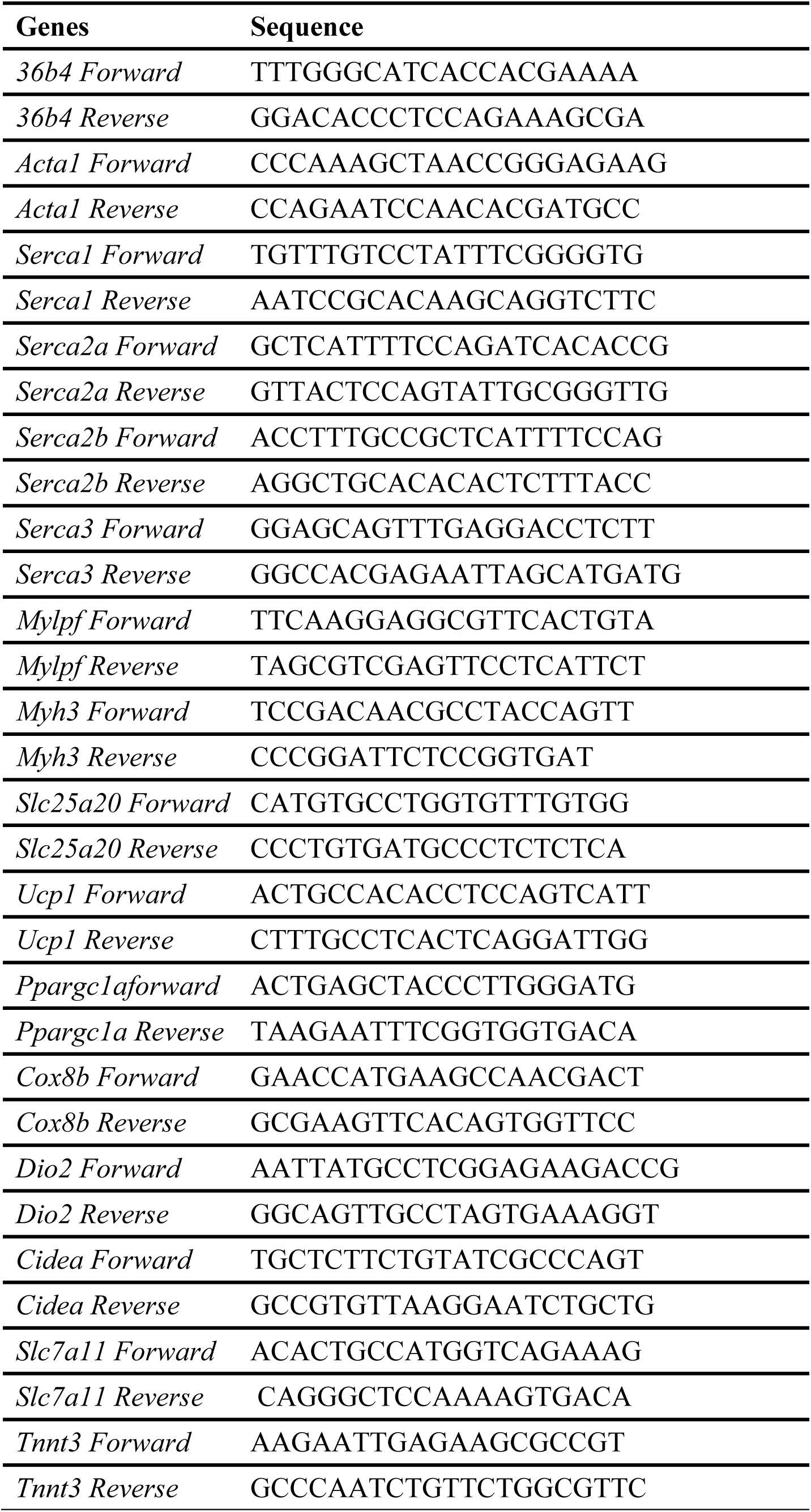
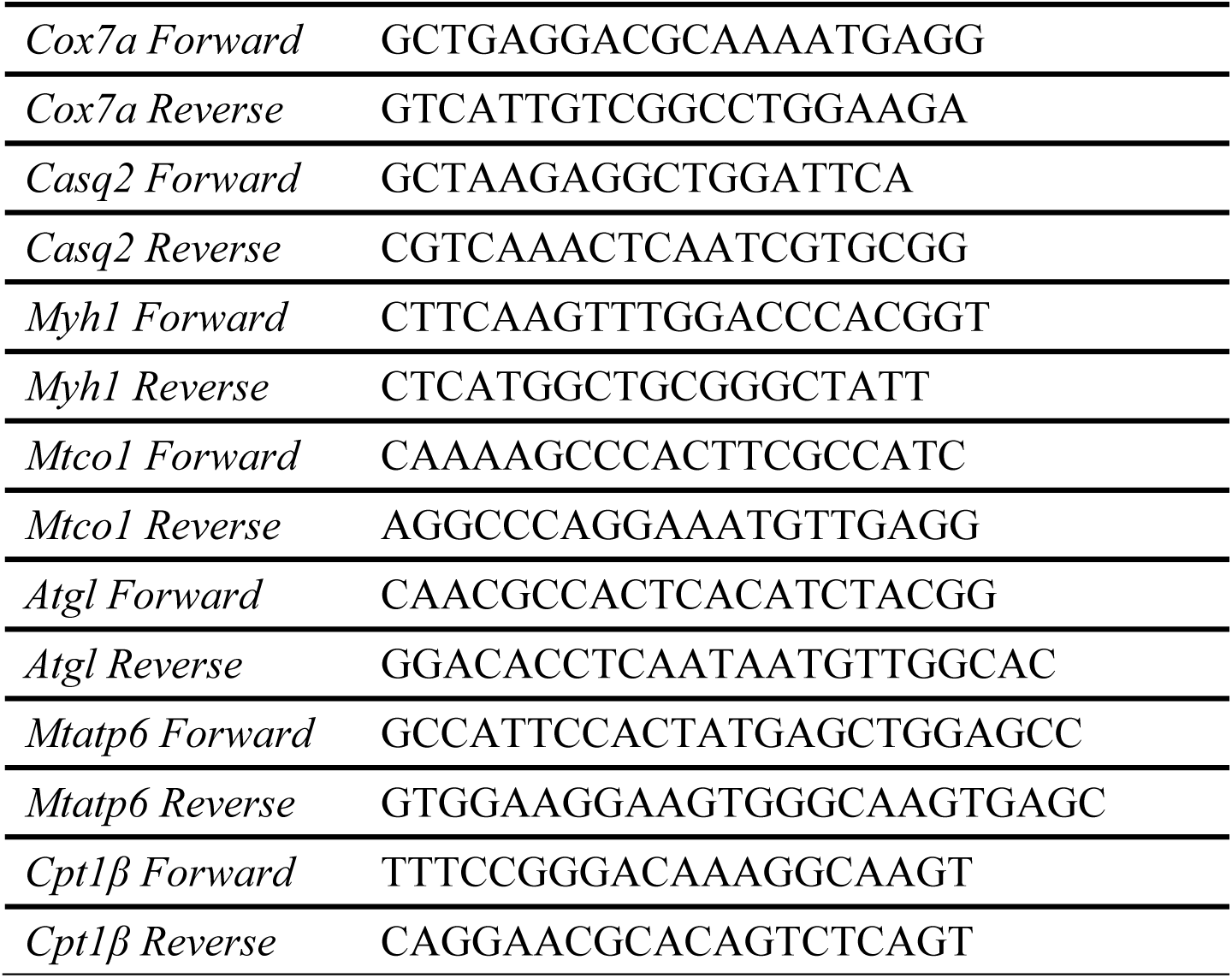
Primer sequences used for RT-qPCR (*Mus musculus*).

**Supplemental Fig. 1.**
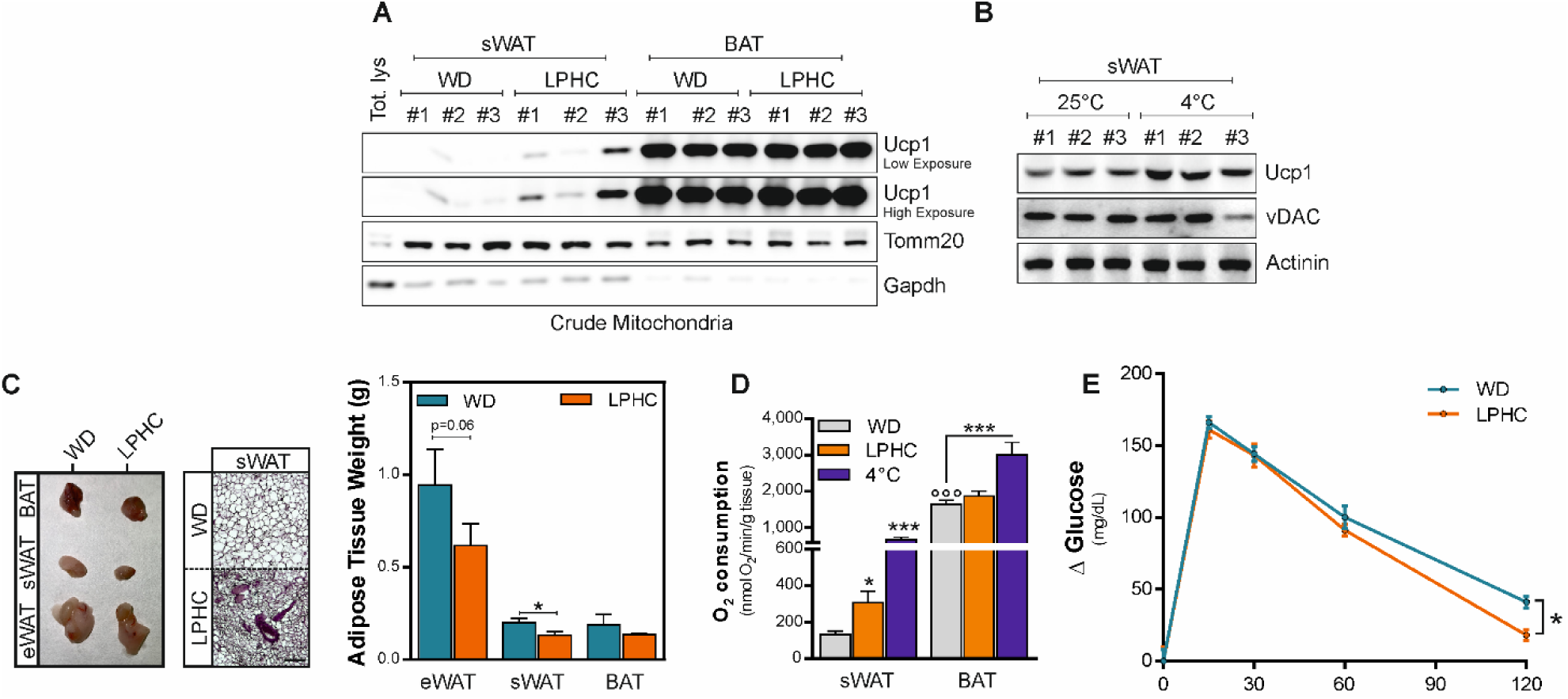
LPHC induces sWAT browning and improves glucose tolerance in mice. **A.** Representative immunoblots of Ucp1 in mitochondria isolated from sWAT of mice fed with WD (n=4 mice) or LPHC diet (n=3 mice) for 2 weeks (A). Tomm20 and Gapdh were used as loading/purity control. **B.** Representative immunoblots of Ucp1 in sWAT of mice exposed to room (25 °C, n=4 mice) or cool temperature (4 °C, n=5 mice) for 24 h. vDAC and Actinin were used as loading controls. **C.** Representative photograph of adipose tissues (left panel) and histochemistry (central panel), and adipose tissues weight (right panel) isolated from mice fed with WD (n=4 mice) or LPHC diet (n=3 mice) for 2 weeks. Bar graphs are presented as mean ± S.D. *p<0.05 LPHC *vs* WD (Student t-test). **D.** Basal oxygen consumption was measured by polarography in sWAT and BAT from mice fed with WD (n=4 mice) or LPHC diet (n=3 mice) for 2 weeks or exposed to cool temperature (4 °C, n=5 mice) for 24 h. Bar graphs are presented as mean ± S.D. *p<0.05, ***p<0.001 LPHC or 4 °C *vs* WD; °°°p<0.001 BAT WD *vs* sWAT WD (one-way ANOVA followed by Dunnett’s multiple comparisons test). **E.** Oral glucose tolerance test was performed in mice fed with WD (n=4 mice) or LPHC diet (n=3 mice) for 2 weeks. Data are presented as mean ± S.D. *p<0.05, LPHC *vs* WD (Student t-test).

**Supplemental Fig. 2.**
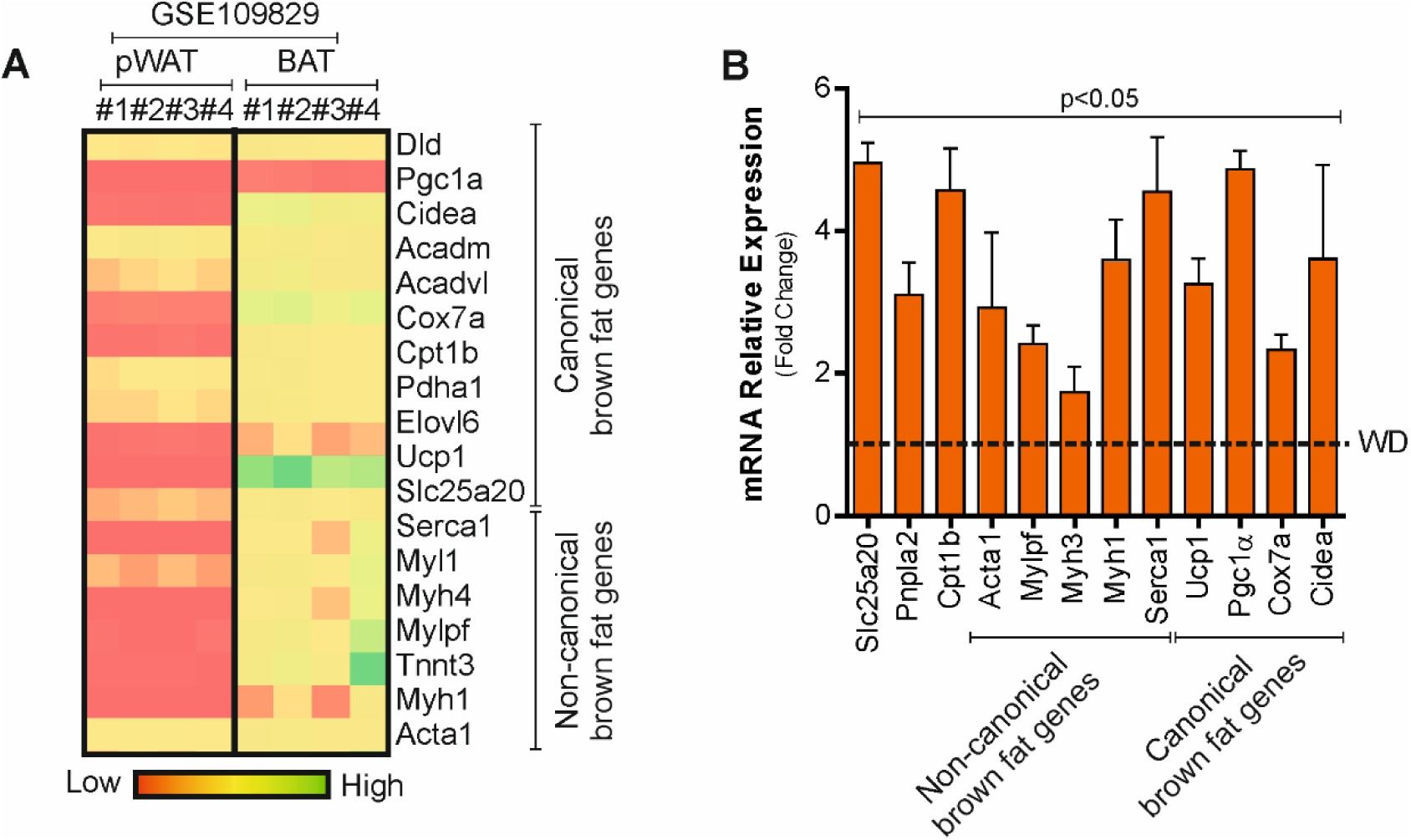
LPHC induces canonical and non-canonical brown fat genes in sWAT. **A.** Comparative expression level of canonical and non-canonical (muscular) brown fat genes in perigonadal white (pWAT) or BAT (GSE109829). Heatmap describes raw expression values. **B.** Single gene expression analysis of canonical and non-canonical (muscular) brown fat genes in sWAT from mice fed with WD (n=4 mice) or LPHC diet (n=3 mice) for 2 weeks. Data are presented as mean ± S.D. p<0.05, LPHC *vs* WD (Student t-test).

**Supplemental Fig. 3.**
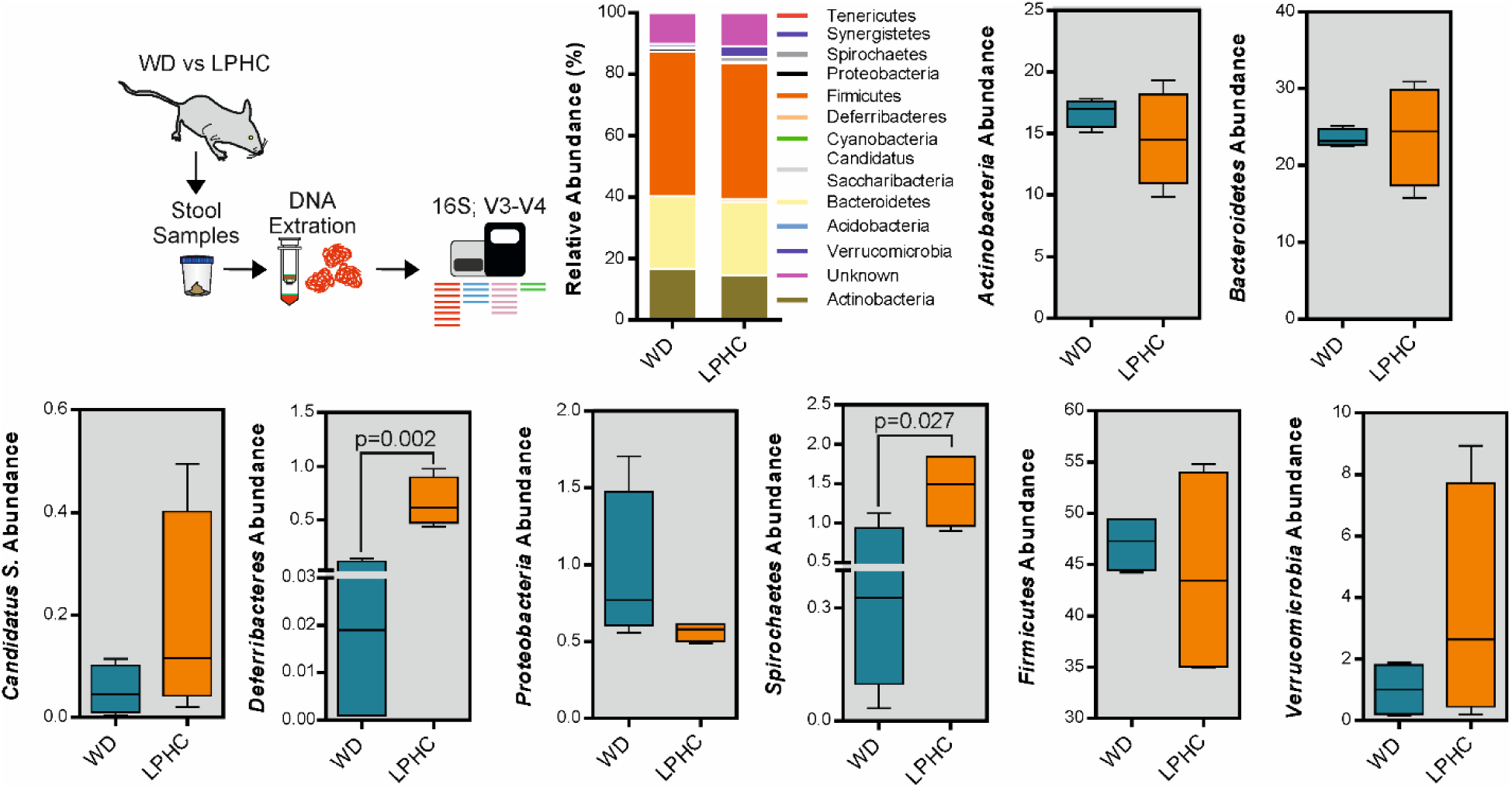
Microbiota reshaping during LPHC diet. Schematic representation of experimental plan and percentage of each bacteria sequence in all sequence reads. Box plots are representative Phyla Operational Taxonomic Unit (OTU) abundance (%) of feces of WD (n=4 mice) and LPHC fed mice (n=4 mice). Data are presented as min to max (Student t test).

**Supplemental Fig. 4.**
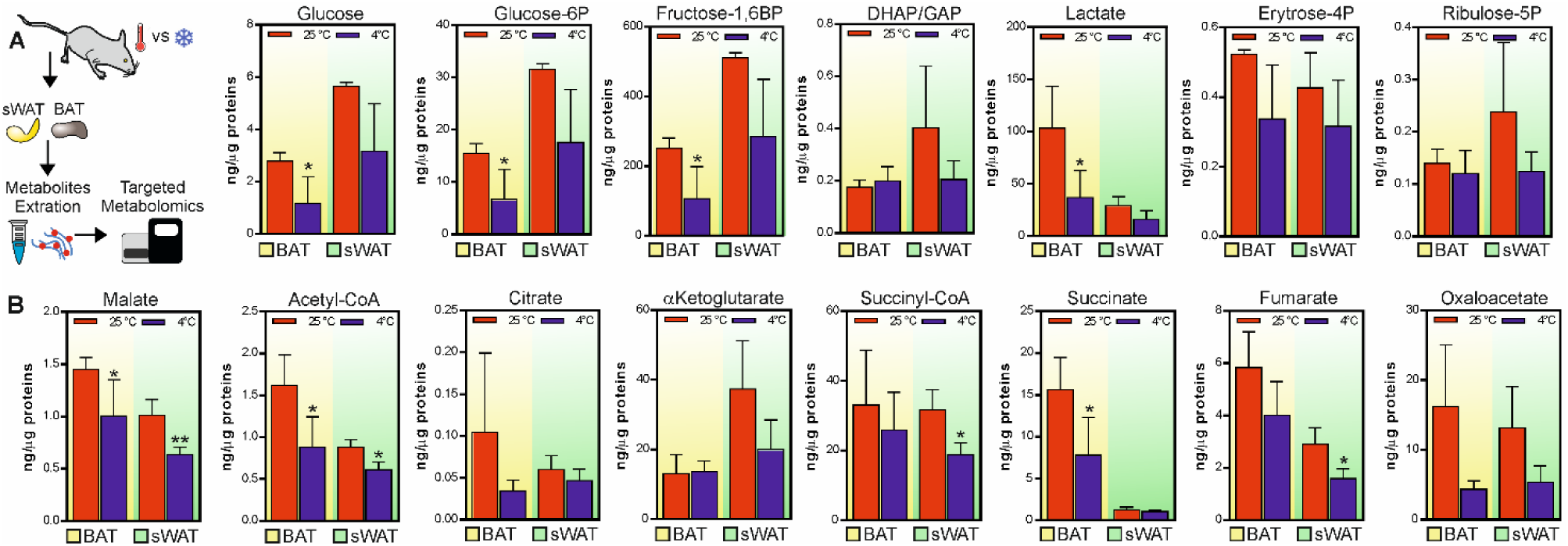
Cold exposure moves glycolysis and TCA cycle targeting metabolites mainly in BAT A, B. Schematic representation of experimental plan; metabolites targeting glycolysis (A) and TCA cycle (B) in BAT (yellow area) and sWAT (green area) from mice exposed to room (25 °C, n=4 mice) or cool temperature (4 °C, n=5 mice) for 24 h. Data are presented as mean ± S.D. *p<0.05, **p<0.01 4 °C *vs* 25 °C (Student t-test).

**S5.**
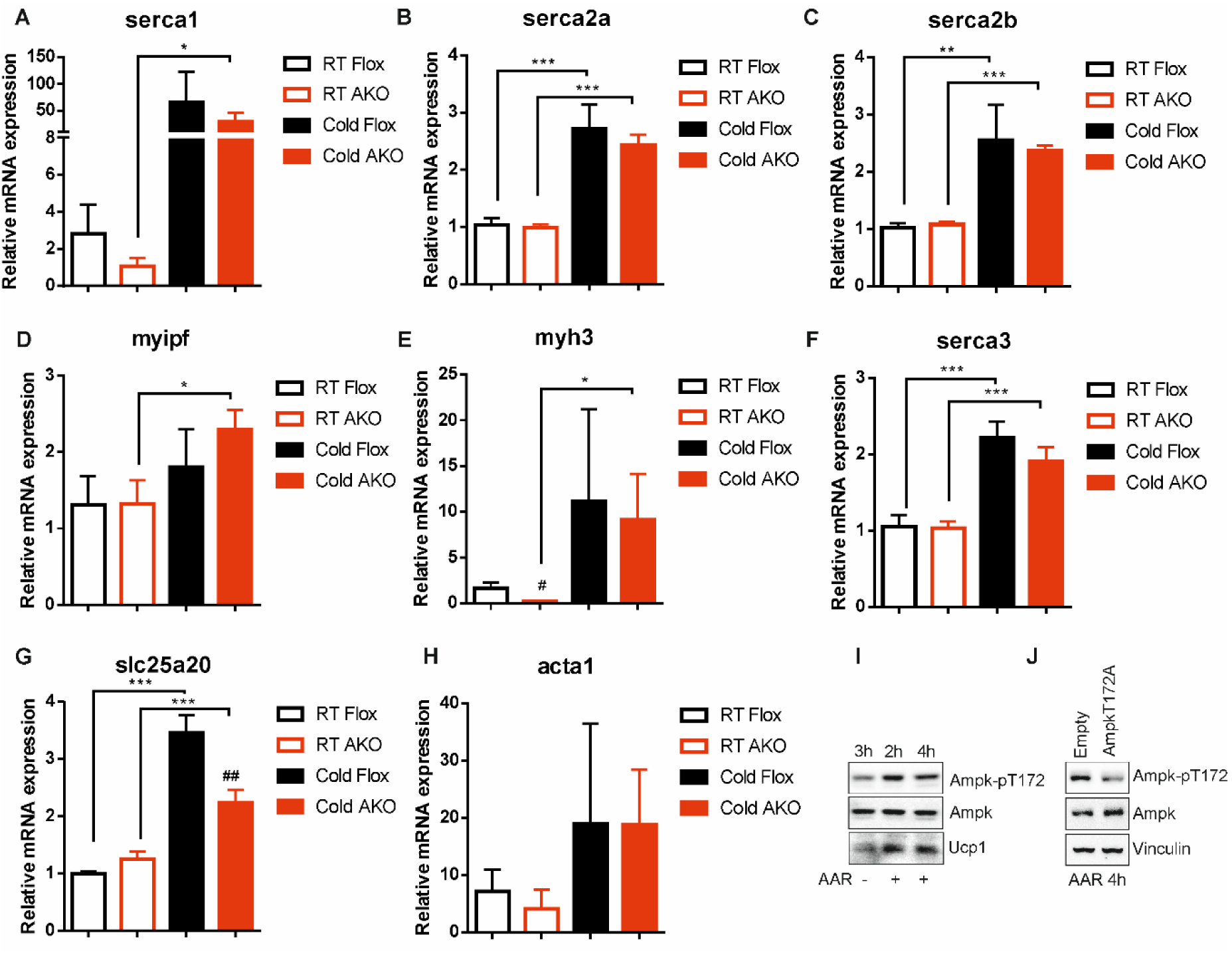
BAT of AKO mice showed unaffected expression levels of muscular genes. **A-H.** Single gene expression analysis of muscular genes in BAT from Flox or AKO mice exposed to room (22 °C) or cool temperature (4 °C) for 48 h. Data are presented as mean ± S.E.M. *p<0.05, **p<0.01, ***p<0.001 4°C *vs* 22°C; #p<0.05, ##p<0.01, AKO 4 °C *vs* Flox 4 °C (one-way ANOVA followed by Dunnett’s multiple comparisons test). **I.** Representative immunoblots of AMPK (basal and phosphorylated) and Ucp1 in X9 beige adipocytes cultured in a medium poor in amino acids (AAR). Basal AMPK was used as loading control. **J.** Representative immunoblots of basal and phosphorylated AMPK in WT and AMPKT172A X9 beige adipocytes cultured in a medium poor in amino acids (AAR). Vinculin was used as loading control.

